# Phospholipids are imported into mitochondria by VDAC, a dimeric beta barrel scramblase

**DOI:** 10.1101/2022.10.17.512472

**Authors:** Helene Jahn, Ladislav Bartoš, Grace I. Dearden, Jeremy S. Dittman, Joost C. M. Holthuis, Robert Vácha, Anant K. Menon

## Abstract

Mitochondria are double-membrane-bounded organelles that depend critically on phospholipids supplied by the endoplasmic reticulum. These lipids must cross the outer membrane to support mitochondrial function, but how they do this is unclear. We identified the voltage-dependent ion channel (VDAC), an abundant outer membrane protein, as a scramblase-type lipid transporter that catalyzes lipid entry. On reconstitution into membrane vesicles, dimers of human VDAC1 and VDAC2 catalyze rapid transbilayer translocation of phospholipids by a mechanism that is unrelated to their channel activity. Coarse-grained molecular dynamics simulations of VDAC1 reveal that lipid scrambling occurs at a specific dimer interface where polar residues induce large water defects and bilayer thinning. The rate of phospholipid import into yeast mitochondria is an order of magnitude lower in the absence of VDAC homologs, indicating that VDACs provide the main pathway for lipid entry. Thus, VDAC isoforms, members of a superfamily of beta barrel proteins, moonlight as a new class of phospholipid scramblases - distinct from alpha-helical scramblase proteins - that act by an unprecedented mechanism to import lipids into mitochondria.

## INTRODUCTION

The double membrane of mitochondria is composed of phospholipids which are supplied by the endoplasmic reticulum (ER) or assembled *in situ* from ER-derived phospholipid precursors (Horvath & Daum, 2013; Tamura *et al*, 2020; Tatsuta & Langer, 2017). For example, cardiolipin, the signature lipid of mitochondria, is synthesized at the matrix side of the inner mitochondrial membrane (IMM) from the ER-derived phospholipid phosphatidic acid (PA), and subsequently remodeled to its mature form in the inter-membrane space (IMS) by enzymatic exchange of acyl chains with ER-derived phosphatidylcholine (PC) (Horvath & Daum, 2013; Schlame & Greenberg, 2017; Tamura *et al*., 2020; Tatsuta & Langer, 2017). This poses a considerable lipid trafficking problem, as, after delivery to the cytoplasmic face of the outer mitochondrial membrane (OMM) by non-vesicular mechanisms (Reinisch & Prinz, 2021), PA, PC, and other phospholipids must cross the barrier of the OMM before moving on to the IMM (Fig. 1A). Previous work suggested that phospholipids are scrambled across the OMM, i.e., flip-flopped across the OMM by an ATP-independent mechanism (Dolis *et al*, 1996; Janssen *et al*, 1999; Jasinska *et al*, 1993; Lampl *et al*, 1994; Pomorski & Menon, 2016), but the identity of the OMM scramblase(s) is not known.

**Figure 1.**
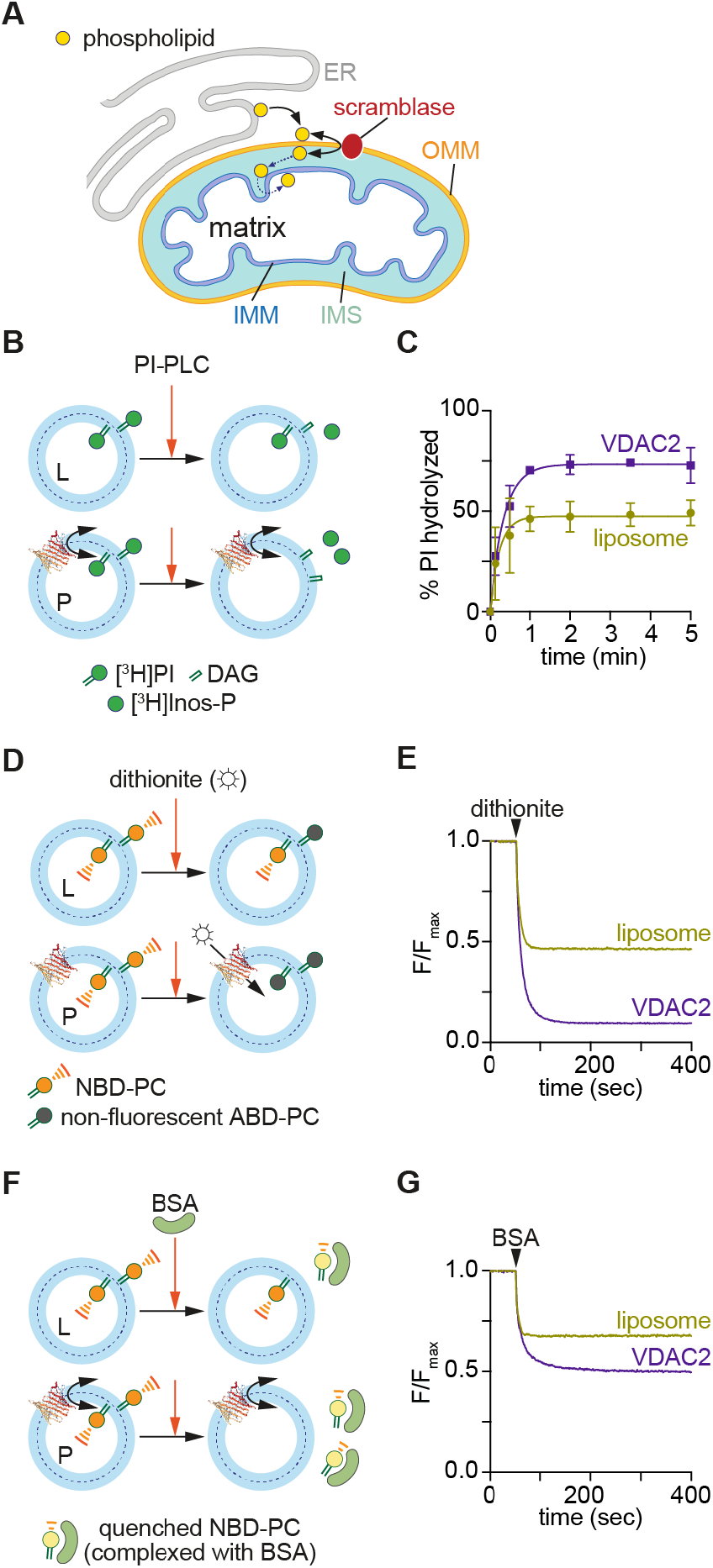
VDAC2 is a phospholipid scramblase. **A.** Schematic showing phospholipid transport from the ER to all bilayer leaflets of the mitochondrial double membrane. Non-vesicular mechanisms deliver ER-synthesized phospholipids to the cytoplasmic face of the outer mitochondrial membrane (OMM). The lipids are scrambled across the OMM by a lipid transporter (scramblase) before moving through the intermembrane space (IMS) and across the inner mitochondrial membrane (IMM). **B.** Scramblase assay using [^3^H]phosphatidylinositol ([^3^H]PI). Protein-free-liposomes (L) or VDAC2-proteoliposomes (P) are reconstituted with [^3^H]PI and probed with PI-specific phospholipase C (PI-PLC), which hydrolyzes [^3^H]PI to diacylglycerol (DAG) and [^3^H]inositol cyclic phosphate ([^3^H]Inos-P). Only [^3^H]PI molecules in the outer leaflet are hydrolyzed in protein-free-liposomes, whereas all [^3^H]PI will be hydrolyzed in scramblase-containing vesicles where [^3^H]PI from the inner leaflet is scrambled to the outer leaflet. **C.** Time-course of PI-PLC-mediated hydrolysis of [^3^H]PI in protein-free-vesicles (liposome) versus vesicles reconstituted with VDAC2 at a theoretical protein/vesicle copy number of 30. Data points are shown as mean and 95% CI, n = 4, and fitted to a single exponential function. **D.** Channel assay using NBD-PC. Vesicles are reconstituted as in panel B, but with fluorescent NBD-PC instead of [^3^H]PI. The membrane-impermeant reductant dithionite bleaches approximately 50% of the NBD-PC fluorescence in protein-free liposomes (L), corresponding to those molecules located in the outer leaflet. For vesicles containing a VDAC2 channel, all NBD-PC molecules are expected to be bleached as dithionite can enter the vesicles. **E.** Time-course of NBD-PC fluorescence (normalized to the starting value) in liposomes and VDAC2-vesicles on adding dithionite. **F.** Scramblase assay using NBD-PC. Vesicles are reconstituted as in panel D and treated with fatty acid-free BSA which extracts NBD-PC from the outer leaflet. In complex with BSA, NBD-PC fluorescence is partly quenched, ∼60% lower than when it is in the membrane. For protein-free liposomes, fluorescence is expected to drop by ∼30%, whereas for scramblase-containing vesicles fluorescence is expected to drop by ∼60%. **G.** Time-course of NBD-PC fluorescence (normalized to the starting value) in liposomes and VDAC2-vesicles on adding BSA.

The Voltage Dependent Anion Channel (VDAC) is an abundant, multi-functional OMM β-barrel protein with three isoforms (VDAC1-3) in mammals and two (Por1-2) in budding yeast (Bergdoll *et al*, 2017; De Pinto, 2021; Di Rosa *et al*, 2021; Raghavan *et al*, 2012; Shoshan-Barmatz *et al*, 2015; Zeth & Zachariae, 2018). It forms the pore through which ATP and other metabolites cross the OMM and plays a key role in apoptosis (Bergdoll *et al*., 2017; De Pinto, 2021; Di Rosa *et al*., 2021; Raghavan *et al*., 2012; Shoshan-Barmatz *et al*., 2015; Zeth & Zachariae, 2018). The different VDAC isoforms also play non-redundant roles in cells (Bergdoll *et al*., 2017; Messina *et al*, 2012; Raghavan *et al*., 2012). Previous molecular dynamics (MD) simulations of mouse VDAC1 (Villinger *et al*, 2010) indicated membrane thinning near an outward-facing glutamate residue (E73), situated midway along the β-barrel. This thinning was observed to promote phospholipid flip-flop (Villinger *et al*., 2010), with headgroups transiently approaching E73 as the lipids crossed from one side of the bilayer to the other. Using coarse-grained MD (CGMD) simulations, we confirmed that the membrane-facing surface of VDAC1 provides a low energy path for phospholipid transit across the bilayer, populated by polar residues and centered on E73 (Fig. S1). These observations suggested that VDAC1 -and perhaps other VDAC isoforms - may be able to facilitate rapid phospholipid flip-flop, thereby providing the scramblase activity needed to move lipids across the OMM for mitochondrial function.

Here we show that human VDAC1 and VDAC2, as well as their yeast ortholog Por1, are phospholipid scramblases: on reconstitution into synthetic vesicles these proteins catalyze rapid transbilayer translocation of phospholipids by a mechanism that is unrelated to their channel activity. We also show that VDACs represent the main mechanism by which phospholipids cross the OMM as the absence of VDAC homologs in yeast mitochondria leads to an order-of-magnitude reduction in the transport rate. However, the originally hypothesized transport pathway centered on residue E73 appears not to play an important role. Biochemical studies and CGMD simulations indicate that fast scrambling requires VDAC dimerization, such that phospholipids transit between the leaflets of the bilayer at a specific dimer interface where polar residues induce large water defects and bilayer thinning.

## RESULTS

### VDAC2 scrambles phospholipids in reconstituted vesicles

To test if human VDAC proteins have scramblase activity we used a previously described assay (Wang *et al*, 2018)(Fig. 1B). We purified correctly folded human VDAC2 after producing it recombinantly in *E. coli* (Fig. S2) and reconstituted it into large unilamellar vesicles composed mainly of PC with a trace quantity of [_3_H]inositol-labeled phosphatidylinositol ([_3_H]PI). Protein-free vesicles were prepared in parallel. On adding bacterial PI-specific phospholipase C (PI-PLC) to the protein-free vesicles we observed rapid (t_1/2_∼10 s) hydrolysis of ∼50% of the [^3^H]PI, corresponding to those molecules residing in the outer leaflet of the vesicles and therefore accessible to the enzyme. However, in samples reconstituted with VDAC2, the extent of hydrolysis increased to ∼75% (Fig. 1C) indicating flipping of [_3_H]PI molecules from the inner to the outer leaflet in a large fraction of the vesicles, at a rate (t_1/2_∼14 s) comparable to that of the hydrolysis reaction. As flipping was observed in the absence of any metabolic energy supplied to the system, this result indicates that VDAC2 is a scramblase (Fig. 1C).

We further characterized lipid scrambling by VDAC2 using fluorescence-based methods (Chang *et al*, 2004; Kubelt *et al*, 2002; Menon *et al*, 2011). VDAC2 was reconstituted into vesicles containing a trace quantity of 1-myristoyl-2-C6-NBD-PC, a fluorescent PC analog. As VDAC2 is a channel/pore, reconstitution efficiency was assessed using dithionite-mediated bleaching of NBD-PC (Fig. 1D). All NBD-PC molecules are expected to be bleached in a vesicle that contains a VDAC2 channel, which permits dithionite entry, whereas only those molecules in the outer leaflet will be bleached in vesicles lacking a channel. Fig. 1E shows that >90% of the NBD-PC is bleached, indicating efficient reconstitution of VDAC2 (in comparison, ∼53% bleaching was observed in protein-free vesicles, as expected for 150 nm-diameter vesicles). Next, scramblase activity was assayed using fatty acid-free bovine serum albumin (BSA), which effectively extracts NBD-PC molecules from the outer leaflet of the vesicles, resulting in a decrease in fluorescence as the quantum efficiency of NBD-PC complexed to BSA is ∼60% lower than that of NBD-PC in the membrane (Chang *et al*., 2004; Kubelt *et al*., 2002; Menon *et al*., 2011)(Fig. 1F). Thus, quantitative extraction of outer leaflet NBD-PC from protein-free liposomes causes ∼30% reduction in fluorescence (Fig. 1G). For vesicles reconstituted with VDAC2, the observed fluorescence reduction is ∼55% (Fig. 1G), the extent predicted if ∼90% of the vesicles include a scramblase, consistent with the reconstitution efficiency deduced from the channel assay. Curve fitting revealed that fluorescence loss on BSA treatment of VDAC2-containing vesicles is characterized by a rapid phase (t_1/2_∼5-10 s, also seen in protein-free samples) corresponding to the extraction of NBD-PC located initially in the outer leaflet, followed by a slower phase (t_1/2_∼50 s) which we attribute to trans-bilayer movement of NBD-PC.

To test whether the ability to scramble lipids is unique to the VDAC β-barrel, we purified an unrelated β-barrel protein, Pet464 (Fig. S3A), the 12-stranded β-barrel portion of the *E. coli* Pet autotransporter (Barnard *et al*, 2007; Yuan *et al*, 2018). Vesicles reconstituted with Pet464 had channel activity as expected, albeit with a lower dithionite permeation rate compared with VDAC2 (Fig. S3B), but no scramblase activity (Fig. S3C). Consistent with these data, previous work showed that OmpT, a 10-stranded bacterial β-barrel also lacks scramblase activity (Kol *et al*, 2003). These results indicate that scrambling is not a general property of β-barrel proteins but rather a specific property of VDAC2. We conclude that VDAC2 is a scramblase, capable of transporting both anionic (PI) and zwitterionic (PC) phospholipids.

### VDAC dimers support fast scrambling

To investigate the mechanism of VDAC2-mediated phospholipid scrambling we tested the effect of disrupting the predicted lipid transit pathway (Fig. S1) by substituting the membrane-facing glutamate residue (E84 in human VDAC2) with leucine. We found, unexpectedly, that the E84L mutation had no detectable impact on the ability of VDAC2 to scramble NBD-PC on the timescale of our experiment (Fig. S4), suggesting that the hypothesized pathway is not the principal avenue for lipid scrambling. We therefore considered the alternative possibility that scramblase activity may depend on VDAC’s quaternary structure.

VDAC dimers and oligomers have been suggested to be physiologically important, for example in apoptosis, and their formation is regulated by various factors, including pH (Bergdoll *et al*, 2018; Betaneli *et al*, 2012; Keinan *et al*, 2013; Kim *et al*, 2019; Schredelseker *et al*, 2014; Shoshan-Barmatz *et al*, 2013; Zalk *et al*, 2005). These oligomeric states have been visualized in the OMM by atomic force microscopy (Goncalves *et al*, 2008; Hoogenboom *et al*, 2007) and cryoelectron microscopy (Leung *et al*, 2021; Mannella, 1982), as well as in detergent solution and after reconstitution into vesicles (Bayrhuber *et al*, 2008; Bergdoll *et al*., 2018; Betaneli *et al*., 2012; Geula *et al*, 2012; Schredelseker *et al*., 2014; Shoshan-Barmatz *et al*., 2013; Zalk *et al*., 2005). To establish the functional oligomeric state of VDAC in our reconstituted system, we determined vesicle occupancy statistics (Cliff *et al*, 2020; Goren *et al*, 2014; Ploier *et al*, 2016; Stockbridge, 2021). We varied the protein/phospholipid ratio (PPR) of the preparation by reconstituting different amounts of VDAC2 into a fixed quantity of lipid vesicles and used the channel assay to determine the fraction of vesicles, at each ratio, that had been functionalized with a VDAC2 channel. The same analyses were also performed using human VDAC1, which we similarly produced in *E. coli* (Fig. S2). As shown in Fig. 2A,B, both VDAC2 and VDAC1 readily functionalize vesicles with channels, with Poisson analysis indicating that they reconstitute as higher-order structures that we estimate to be decamers (Fig. 2C). This could be because the proteins are intrinsically multimeric as purified and/or multimerize during detergent removal *en route* to reconstitution (Betaneli *et al*., 2012).

**Figure 2.**
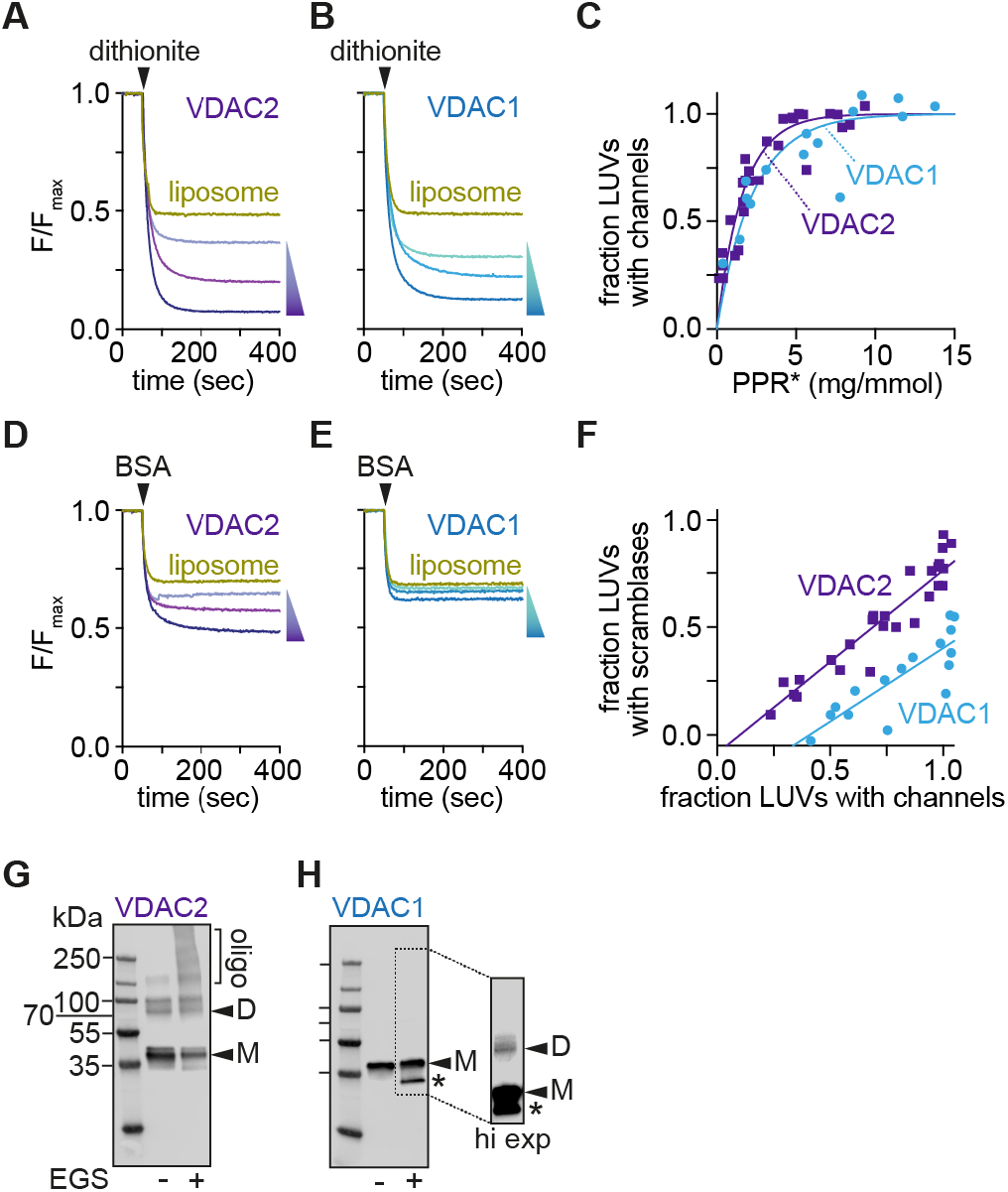
Native VDAC1 scrambles phospholipids poorly. **A, B.** Representative traces corresponding to channel assays performed on vesicles reconstituted with different amounts of VDAC2 (panel A) or VDAC1 (panel B) (Shown are protein concentrations corresponding to theoretical protein/vesicle copy number of 30, 10 and 2). **C.** Protein-dependence plot showing functionalization of vesicles with channel activity, i.e., fraction of large unilamellar vesicles (LUVs) with at least one channel. PPR*, PPR (protein/phospholipid ratio (mg/mmol)), corrected to eliminate the contribution of empty vesicles (Ploier *et al*., 2016). The data were analyzed according to a Poisson statistics model for reconstitution of proteins into individual vesicles. A similar mono-exponential fit constant was obtained for both proteins (∼2 mg/mmol). **D, E.** Representative traces corresponding to scramblase assays performed on vesicles reconstituted with different amounts of VDAC2 (panel D) or VDAC1 (panel E) as in panel A and B. F. Correlation between the fraction of LUVs showing channel activity (dithionite assay) and vesicles with scramblase activity (BSA-back extraction assay). Data points correspond to LUVs reconstituted with different amounts of protein. **G, H.** Crosslinking of VDAC proteins after reconstitution into vesicles. Reconstituted LUVs were treated with EGS to crosslink proteins in proximity. The samples were analyzed by SDS-PAGE immunoblotting using antibodies against the N-terminal His tag. Reconstituted VDAC2 shows a significant population of dimers/multimers (panel G) whereas reconstituted VDAC1 is predominantly monomeric (panel H, side panel shows a brighter signal to enable visualization of a faint dimer band).

The same vesicle preparations were next taken for scramblase assays. VDAC2 showed robust scramblase activity as expected, the fraction of functionalized vesicles scaling with the PPR as seen for the channel assays (Fig. 2D). However, VDAC1 was surprisingly less potent (Fig. 2E). To evaluate this difference quantitatively, we graphed the fraction of scramblase-active vesicles versus the fraction of channel-active vesicles for both isoforms (Fig. 2F). For VDAC2, these fractions were highly correlated, with ∼85% of the channel active vesicles displaying scramblase activity over the range of the PPRs that we tested. However, for VDAC1, scramblase activity was essentially undetectable on the timescale of the assay until sufficient protein was reconstituted to populate about a third of the vesicles with channels. As the fraction of channel-active vesicles increased beyond ∼0.3, the fraction of scramblase-active vesicles also increased but less efficiently (correlation ∼70%) than in the case of VDAC2 (Fig. 2F).

To understand the molecular basis for the difference in the behavior of these otherwise highly similar isoforms of VDAC (75% identity, 91% positive match (Messina *et al*., 2012; Schredelseker *et al*., 2014)), we used chemical crosslinking to probe their quaternary structure after reconstitution. VDAC1-and VDAC2-containing vesicles were treated with EGS, an amine-reactive crosslinker which has been previously used to determine the oligomeric nature of VDAC (Geula *et al*., 2012; Zalk *et al*., 2005)(Fig. S5A). Immunoblotting revealed that in the absence of the crosslinker, both proteins were detected mainly as monomers, with a small fraction of VDAC2 forming SDS-resistant dimers. However, the crosslinker efficiently captured dimers and multimers of VDAC2 (Fig. 2G) whereas VDAC1 was recovered mainly in monomeric form (Fig. 2H). We infer that despite their similar incorporation into vesicles as multimers (Fig. 2C), VDAC2 molecules remain associated in the membrane such that complexes can be captured by EGS crosslinking, whereas VDAC1 molecules dissociate, thereby losing their ability to facilitate fast scrambling (assays run over an extended time frame reveal that VDAC1 scrambles lipids slowly, with a long half-time of ∼4 h (Fig. S6)). These data suggest that a dimer or higher order multimer of VDAC is necessary for rapid scrambling.

Covalent VDAC1 dimers can be formed by crosslinking in detergent (Fig. S7) providing an opportunity to determine whether dimerization is sufficient for scrambling. We identified conditions in which we could crosslink VDAC1 efficiently with EGS after purifying it in LDAO detergent (Fig. S7 and Fig. 3A). Immunoblot analysis revealed that the crosslinked sample, VDAC1^x^, consisted principally of dimers, with a small fraction of higher order multimers (Fig. 3A). The EGS-captured dimers likely correspond to VDAC1 protomers oriented in parallel, interacting via strands β1,2,18,19 (Fig. S5C,D), similar to the common dimeric structure reported for human and rat VDAC1 (Bayrhuber *et al*., 2008; Geula *et al*., 2012), and zebrafish VDAC2 (Schredelseker *et al*., 2014). Importantly, the parallel orientation of the VDAC1 protomers within the dimer structure is supported by spectroscopic analyses of LDAO-solubilized mouse VDAC1 (Bergdoll *et al*., 2018) and structural studies of human VDAC1 purified in LDAO (Bayrhuber *et al*., 2008).

**Figure 3.**
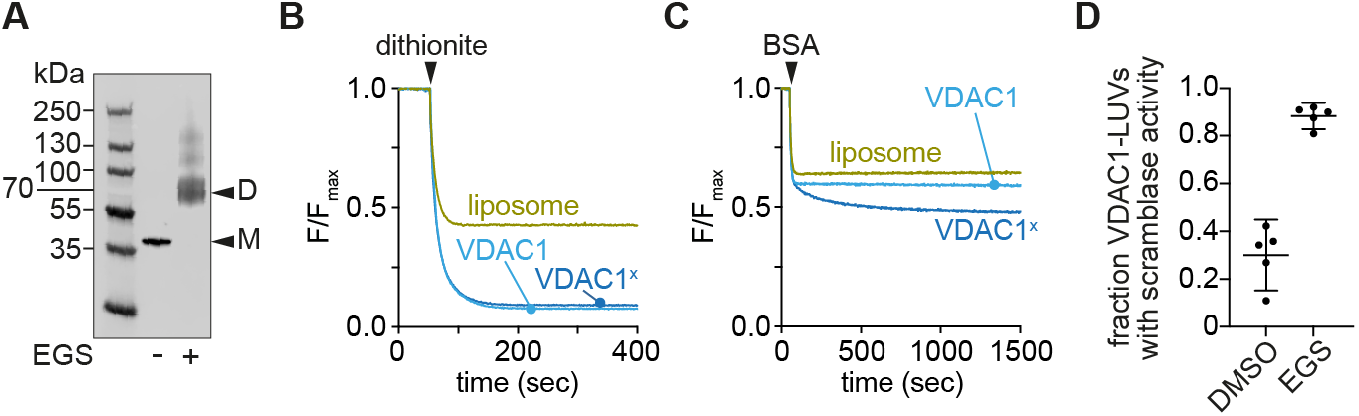
Crosslinked VDAC1 dimers have phospholipid scramblase activity. **A.** VDAC1 in LDAO was cross-linked using EGS, or mocked treated with DMSO, and visualized by SDS-PAGE immunoblotting using antibodies against the N-terminal His tag. **B,C.** Representative dithionite (B) and BSA-back extraction (C) traces with cross-linked (VDAC1^x^) or mock treated (VDAC1) samples reconstituted into LUVs at a theoretical copy number of 30 proteins/vesicle. Normalized fluorescence traces are shown. **D.** Fraction of VDAC1-vesicles with scramblase activity (reconstituted at a theoretical protein/vesicle copy number of 30). Data points shown as individual points, as well as the mean ± 95% CI, n = 5, p <0.0001. After EGS cross-linking, nearly all vesicles that show channel activity also have scramblase activity.

Reconstitution of VDAC1^x^ and mock-treated VDAC1 showed that they both functionalized vesicles efficiently as evinced by channel activity (Figure 3B). However, whereas the mock-treated protein was barely active as a scramblase, VDAC1^x^ showed robust scrambling activity when reconstituted at the same PPR (Fig. 3C). Quantification of the results (Fig. 3D) showed that ∼90% of channel-active vesicles populated with VDAC1^x^ are also scramblase active, whereas this number is only ∼30% in the case of reconstituted VDAC monomers. We note that VDAC1^x^ scrambles lipids ∼2-fold more slowly than native dimers/multimers of VDAC2 (Fig. 3C vs 1G), perhaps because of the physical constraints imposed by crosslinking.

To extend our results with EGS, we used DSP, a cleavable crosslinker containing a dithiothreitol (DTT)-susceptible disulfide bond (Fig. S5B). On crosslinking LDAO-solubilized VDAC1 with DSP, we obtained mainly dimers (Fig. S5E), which could be restored to monomers upon DTT treatment (Fig. S5E). DSP-crosslinked VDAC1 had scramblase activity (Fig. S5F,G), like VDAC^x^, which was largely lost on DTT-treatment of the reconstituted sample (Fig. S5F,G). We conclude that VDAC dimerization is both necessary and sufficient to generate an efficient scramblase, and that the dimer is the likely minimal functional oligomeric state of the protein.

### The VDAC pore is not relevant for rapid scrambling

Our reconstitution experiments with VDAC1 indicate that it functionalizes vesicles efficiently with channel activity but does not facilitate scrambling unless it is dimerized by crosslinking prior to reconstitution or reconstituted at a high PPR to promote dimer formation *in situ*. Two useful corollaries emerge from this result. First, monomeric VDAC1 provides a potent negative control for our scramblase assay, reinforcing our result with the Pet464 ý-barrel (Fig. S3C) - thus, mere reconstitution of a β-barrel is not sufficient for rapid scrambling. Second, the inability of monomeric VDAC1 to promote rapid scrambling indicates that the VDAC pore does not participate directly in this process, and that the pathway for transbilayer lipid transit relies on unique features of VDAC’s membrane-facing surface created by dimerization. This latter point is reinforced in experiments noted above where we measured slow scrambling by VDAC1 monomers (Fig. S6). Thus, we prepared monomeric VDAC1-containing vesicles without NBD-PC, and then added the NBD-PC exogenously. Soon after addition, all the NBD-PC could be extracted by albumin, consistent with its deposition in the outer leaflet of the vesicles. However, over a period of 30 h, slow scrambling enabled these lipids to equilibrate with the inner leaflet from which they could no longer be extracted. If NBD-PC molecules could transit through the VDAC1 pore, then they would have been quantitatively extracted at all time points over the incubation period. The presence of a protected (inner leaflet) pool of NBD-PC confirms that the VDAC1 pore is not relevant for scrambling.

### Molecular dynamics simulations reveal a phospholipid transit pathway at a specific dimer interface

To understand how VDAC dimers scramble phospholipids we used coarse-grained molecular dynamics to simulate different dimers. We chose two symmetric dimers, with interfaces mediated by strands β1,2,18,19 (dimer-1) (Fig. 4A,B) as previously reported (Bayrhuber *et al*., 2008; Geula *et al*., 2012; Schredelseker *et al*., 2014), and β14-17 (dimer-3), a novel configuration in which the E73 residue is positioned distal to the interface in each protomer (Fig. S8A). We observed a high scrambling rate for dimer-1 (Fig. 4C,D) over 10 μs of simulation time, with PC molecules moving along the edge of the interface while interacting with both protomers (Fig. 4E,F and Movie S1). In contrast, a VDAC1 monomer supported only slow scrambling, as noted experimentally (Fig. S6), with lipids moving along the originally proposed E73-centered transit path (Fig. S1). Unless restrained, dimer-1 re-oriented during the simulation to dimer-1*, with a symmetric interface mediated by strands β2-4 (Fig. S8B), at which point the rate fell ∼4-fold (Fig. 4C,D). Dimer-3 was largely ineffective (Fig. 4D, Fig. S8C), scrambling lipids at a rate only marginally higher than that seen for two individual VDAC1 monomers. These results suggest that not all dimer interfaces support rapid scrambling. In support of this conclusion, we experimentally identified a VDAC1 mutant (VDAC1-5V) which, although highly multimeric after reconstitution as revealed by EGS crosslinking (Fig. S9A), scrambled lipids poorly (Fig. S9). We conclude that a specific dimer interface is required to promote rapid scrambling.

**Figure 4:**
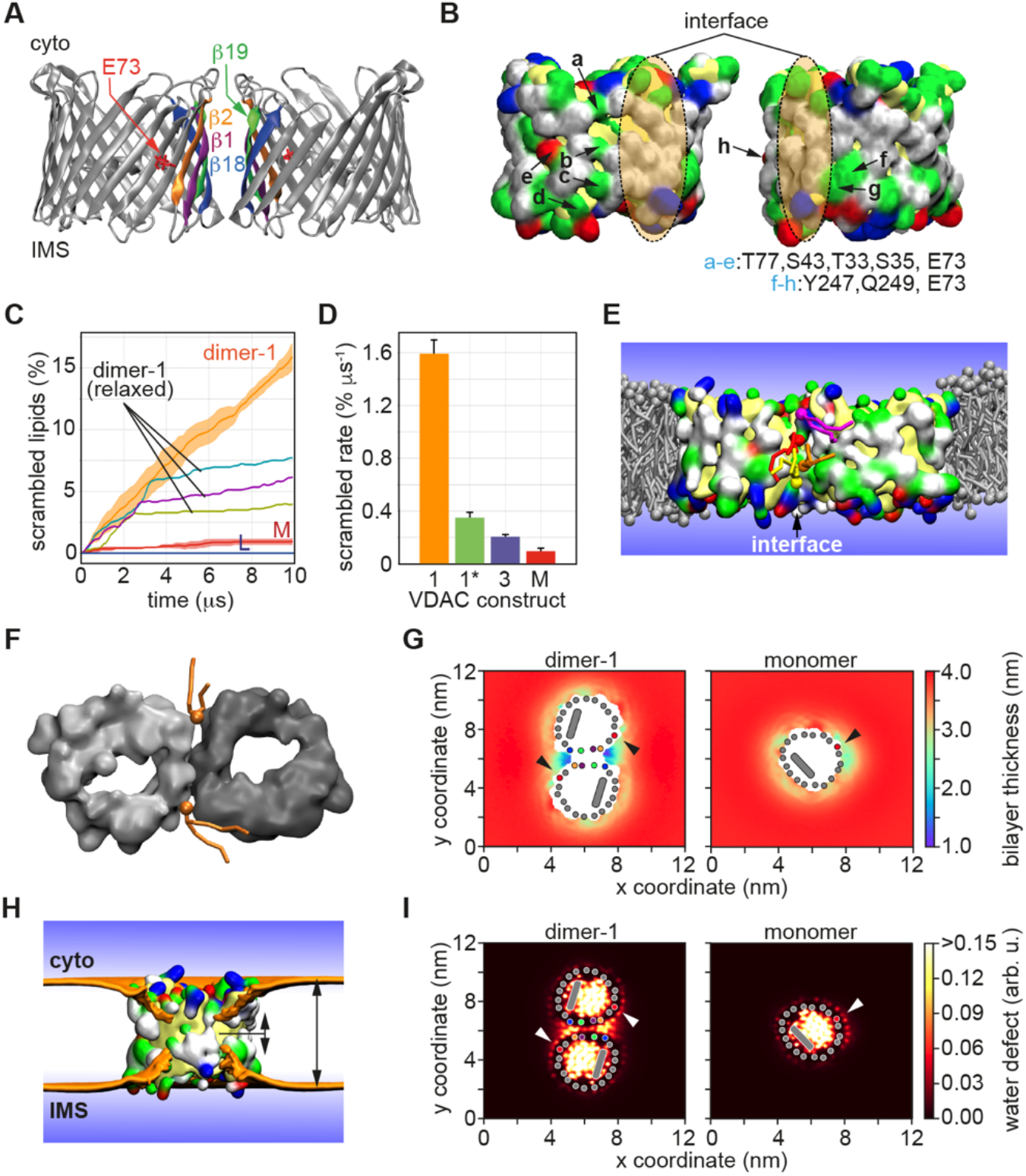
Coarse-grained MD simulations of scrambling by a specific VDAC1 dimer. **A.** Structure of VDAC1 dimer-1. The β-strands at the interface are colored as indicated, and the E73 residue on the β4-strand is marked. Structures of dimer-1* and dimer-3 are shown in Fig. S8. **B.** Surface representation (white = hydrophobic, yellow = backbone, green = hydrophilic, blue = positive, red = negative) showing each protomer, rotated 90° to expose the dimer-1 interface, with the contact region marked with a shaded ellipse; polar residues that flank the interface are indicated according to the key under the structures. The residues with side-chains oriented inwards, i.e., facing the pore of VDAC1, are colored in yellow. **C.** Percentage of lipids scrambled as a function of simulation time. The graphs show the scrambling activity of dimer-1, monomer (M) and protein-free membrane (L) (running average over 200 ns time intervals, shading=68% confidence interval, n=3). Unless constrained during the simulation, dimer-1 reorients to form dimer-1*. Three individual runs of unconstrained dimer-1 are shown (dimer-1 (relaxed), the change in scrambling rate between 2-4 µs coinciding with reorientation. **D.** Bar chart showing scrambling rate for dimer-1, dimer-1*, dimer-3 and monomer (M) (mean ± SD, n=3). **E.** Snapshot of dimer-1 from the simulation showing phospholipids (multiple colors) transiting between bilayer leaflets along the interface. Membrane phospholipids surrounding the dimer are grey. **F.** Top view snapshot of dimer-1 showing representative phospholipids transiting across the bilayer on both sides of the interface. **G.** Bilayer thinning at the dimer-1 interface and near E73 in monomeric VDAC1 (thickness indicated by the color scale shown at right). Top views are shown, β-strands at the interface are indicated as colored dots (same color scheme as in panel A); the red dot indicates the β4-strand where E73 is located (marked by arrowhead); the N-terminal helix is shown as a grey oblong within the VDAC1 pore. **H.** Snapshot side-view of VDAC dimer-1, with one protomer removed, demonstrating membrane thinning at the interface. The ochre surface indicates the average positions of lipid phosphates from both membrane leaflets. The protein is shown in surface representation. I. Water penetration into the membrane (water defect, color scale shown at right) in the vicinity of the dimer-1 interface and monomeric VDAC1, as indicated.

The dimer-1 interface possesses numerous polar residues (Fig. 4B), but these are clearly insufficient to promote fast scrambling by themselves because the monomeric protein scrambles lipids slowly (Fig. 4C,D, Fig. S6). We therefore considered that the individual polar faces must synergize at the interface, possibly in concert with the membrane, to create a low-energy trans-bilayer path for lipids. Indeed, we found that the bilayer was much thinner adjacent to the interface, reaching just over 2 nm (Fig. 4G,H), and exhibited a large degree of water penetration (Fig. 4I). These features could also be seen with dimer-1*, but to a smaller extent, correlating with its weaker scrambling activity (Fig. S8F,G). In contrast, the membrane was only slightly perturbed in the vicinity of dimer-3 (Fig. S8F,G) and the monomeric protein (Fig. 4G,I). We hypothesized that scrambling efficiency would be reduced if the six polar residues in the dimer-1 interface (T77, S43, T33, S35, Y247, Q249 (Fig. 4B)) were to be replaced with valine. We simulated this construct (dimer-1-mutant) and observed that its ability to thin the membrane and promote water permeation was intermediate between that of dimer-3 and dimer-1* (Fig. S8F,G), and that it is a poor scramblase with a scrambling rate comparable to a single monomeric VDAC (Fig. S8D,E). Finally, evaluation of the interactions of the scrambling lipid along the dimer-1 interface (Fig. S10) demonstrated that a substantial portion of the decrease in the flip-flop barrier likely originates from the interaction between the lipid and the residues of VDAC1 along the scrambling pathway. This is readily seen by the increase in the energy barrier for lipid scrambling in simulations (Fig. S10A) in which the attractive dispersion interactions between the lipid and VDAC1 were turned off.

### Phospholipid import into isolated mitochondria is slowed by an order of magnitude in the absence of VDAC

Our reconstitution data and *in silico* analyses clearly show that specific VDAC dimers are efficient scramblases. To quantify the role of VDACs in scrambling phospholipids across the OMM, we turned to the yeast *Saccharomyces cerevisiae*, an organism with VDAC orthologs (Por1 and Por2) that share 70% sequence similarity to human VDAC (Di Rosa *et al*., 2021). We first tested whether yeast VDACs have scramblase activity, and for this we chose to investigate Por1, which is at least 10-fold more abundant than Por2. We over-expressed Twin-Strep-tagged Por1 (Por1-TS) in yeast, and purified it by affinity chromatography, followed by size exclusion (Fig. S11A). Using the methods outlined in Fig. 1D,F, and Fig. 2G, we found that Por1-TS had both channel and scramblase activities (Fig. S11B,C), and that the reconstituted sample contained a significant proportion of dimers (Fig. S11D). These data indicate that Por1, like VDAC2, is sufficiently dimeric to not require prior crosslinking for scramblase activity (we note that a small amount of endogenous Por1 co-purified with Por1-TS (Fig. S11A), consistent with its ability to form dimers). Thus, Por1 is a scramblase.

To test the role of VDACs in scrambling phospholipids across the OMM we assayed the ability of isolated yeast mitochondria to convert exogenously supplied phosphatidylserine (PS) to phosphatidylethanolamine (PE), a multi-step process requiring transport of PS across the OMM to the site of PS decarboxylase (Psd1) in the IMS (Fig. 5A) (Tatsuta & Langer, 2017). We added NBD-PS to wild-type mitochondria and observed a time-dependent conversion to NBD-PE as visualized by thin layer chromatography (Fig. S12A, WT). As expected, no conversion was observed in mitochondria prepared from *psd111* cells (Fig. S12A, *psd1Δ*). The latter result additionally indicates that our preparations are not contaminated with Psd2, a PS decarboxylase localized to the late secretory pathway. We next prepared mitochondria from *por1Δ*, *por2Δ*, and *por111por211* cells, adjusted the preparations to have similar concentrations (confirmed by immunoblotting against the IMM and OMM proteins Psd1 and Tom40, respectively (Fig. S12D)), and measured the rate of conversion of NBD-PS to NBD-PE. Thin layer chromatograms corresponding to the assays are shown in Fig. S12A (lower panels). Quantification of these data yielded time courses (Fig. S12E), and half-times of transport (Fig. S12F). Mitochondria lacking Por1 produced PE at a significantly lower rate than wild-type mitochondria (Fig. S12F), consistent with a role for VDAC in facilitating efficient trans-bilayer movement of PS across the OMM; elimination of Por2, alone or in combination with Por1, did not produce a detectable change in rate consistent with its relatively low abundance. As explained below, the lower rate of PE production in *por111por211* yeast is because the rate of scrambling decreases substantially, such that it becomes lower than the rate of PS decarboxylation.

Because the conversion of NBD-PS to NBD-PE is a multi-step process (Fig. 5A), the rate of NBD-PE production in the assay is only indirectly correlated with the rate at which NBD-PS is transported across the OMM. To tease out how the elimination of Por1/Por2 affects scrambling across the OMM at least two additional contributions to the aggregate kinetics must be considered: deposition of NBD-PS into the OMM outer leaflet and its enzymatic decarboxylation within the IMS by Psd1 (Fig. 5A). As the final decarboxylation step is likely to be rate-limiting, large changes in the rate of transport of NBD-PS across the OMM would be expected to have relatively modest corresponding changes in the rate of NBD-PE production. To quantify the contribution of VDAC-mediated lipid scrambling to the overall process, we developed a simple kinetic framework comprising PS capture, OMM transit, and enzymatic conversion by Psd1, and fit this model to the measured time course of PE production using either wild-type or *por111por211* mitochondria. A minimal scheme required a four-state model accommodating three distinct transitions (Fig. 5A): first, NBD-PS is reversibly deposited onto the OMM surface with kinetics characterized by rate constants k_0_ and k_1_ assuming pseudo-first order kinetics with an excess of available OMM binding capacity (Fig. S12B, C); second, VDAC reversibly transports NBD-PS across the OMM with out®in and in®out rate constants k_2_ and k_3_, respectively; third, NBD-PS is irreversibly decarboxylated to generate NBD-PE with an effective rate constant k_4_. Note that k_4_ incorporates the intrinsic kinetics of the Psd1 enzyme, as well as any contributions from NBD-PS adsorption/desorption kinetics within the IMS, into a single effective rate constant.

**Figure 5.**
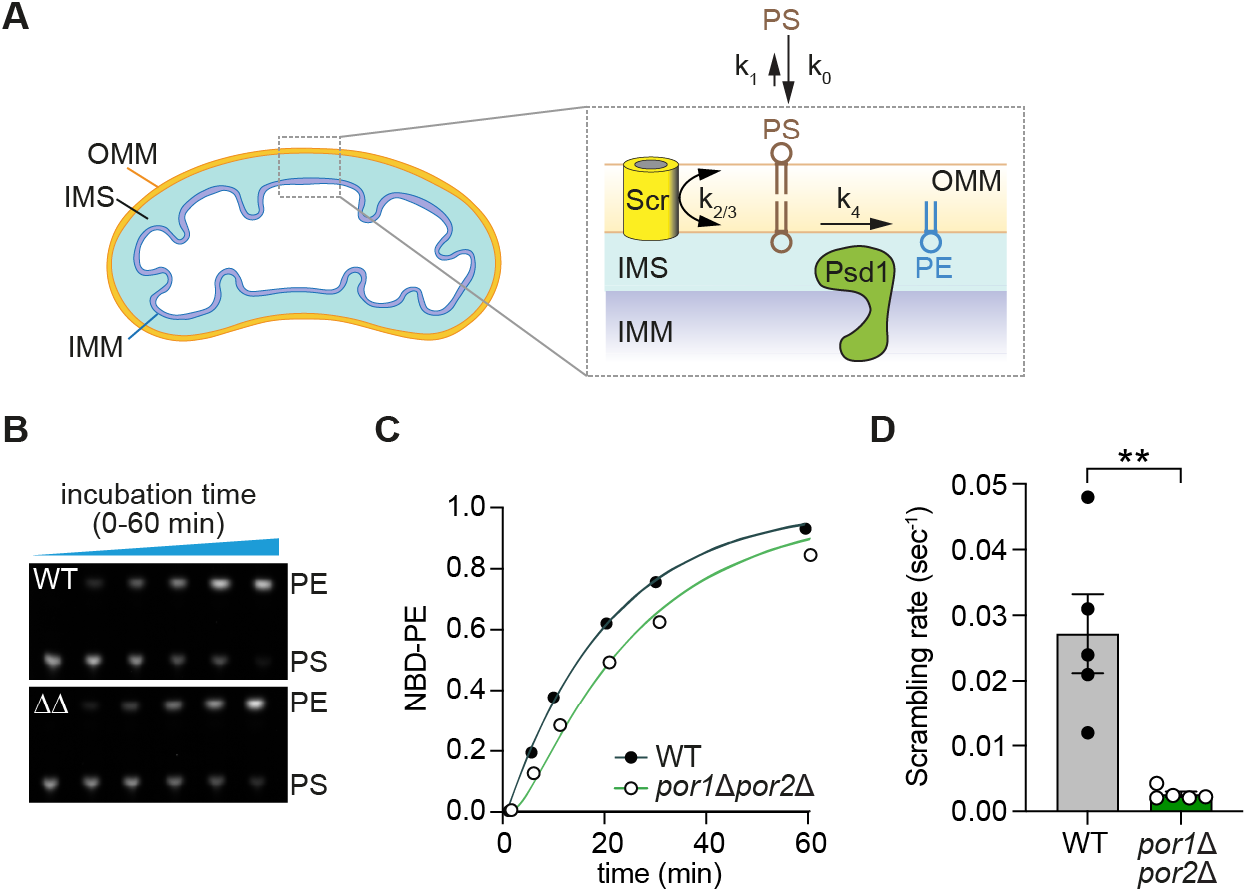
Phospholipid transport across the OMM is slowed more than 10-fold in yeast mitochondria lacking VDAC homologs. **A.** Assay schematic. NBD-PS (indicated PS) is added to yeast mitochondria. After insertion into the outer leaflet of the OMM, NBD-PS flips reversibly across the membrane where it encounters IMM-localized PS decarboxylase (Psd1) which converts it to NBD-PE (indicated PE). Psd1 can act *in trans*, hydrolyzing PS in the inner leaflet of the OMM as shown. It can also act *in cis* on NBD-PS molecules that are delivered to the IMS side of the IMM. The decarboxylation of exogenously supplied NBD-PS can be described using a 4-state kinetic model and 5 effective rate constants as shown: deposition of NBD-PS into the OMM (k_0_), desorption from the OMM (k_1_), scrambling across the OMM (k_2_ and k_3_, presumed to be identical and written as k_2/3_), and irreversible conversion to NBD-PE (k_4_). **B.** Thin layer chromatogram, visualized with a ChemiDoc fluorescence imager, of a decarboxylation assay time-course using mitochondria from wild-type yeast (top panel) and the *por1Δpor2Δ* double mutant (bottom panel, labeled ΔΔ). **C.** Time courses of NBD-PS decarboxylation corresponding to panel B. The data for wild-type mitochondria were analyzed using a 4-state kinetic model (panel A) with k_0_=1 s^-1^, k_1_=0.05 s^-1^, and k_2_=k_3_>0.01 s^-1^ to obtain k_2/3_ and k_4_. The fitting yielded k_4_=0.0018 s^-1^. Data for the *por1Δpor2Δ* double mutant were analyzed using the same kinetic model with k_0_=1 s^-1^, k_1_=0.05 s^-1^ and k_4_=0.0018 s^-1^ to determine k_2/3_. Exemplary fits are shown with circles representing experimental data. **D.** Scrambling rates (k_2/3_) for wild-type and *por1Δpor2Δ* mitochondria obtained as in panel C. The corresponding half-times are 25 and 265 s for wild-type and *por1Δpor2Δ* mitochondria, respectively. ** p < 0.01 (Student’s T test).

We determined the kinetic parameters of the four-state model for wild-type mitochondria as follows. Using liposome-based assays (see Materials and Methods) we determined k_0_ ∼ 1 sec^-1^ and k_1_ ∼ 0.05 sec^-1^ (Fig. S12G-J). Assuming k_2_ = k_3_ > 0.01 sec^-1^ based on previous studies (Janssen *et al*., 1999), and leaving k_4_ unconstrained, we fit the time course of NBD-PE production to obtain k_2_ = k_3_ = 0.027 ± 0.006 sec^-1^ and k_4_ = 0.0018 ± 0.00005 sec^-1^ (Fig. 5B, C). As porin deletion would not be expected to alter k_0_, k_1_, or k_4_, we next fit the kinetic data for the *por111por211* mitochondria to obtain the corresponding values of k_2_ and k_3_. The best-fitting model produced k_2_ = k_3_ = 0.0025 ± 0.0004 sec_-1_, indicating that the rate of lipid scrambling across the OMM decreases by more than 10-fold in the absence of VDAC (Fig. 5D). The loss of >90% of the lipid import capability in mitochondria lacking VDACs indicates that VDAC proteins provide the majority of scramblase activity associated with the OMM.

## DISCUSSION

We identified the ý-barrel proteins human VDAC1 and VDAC2, as well as their yeast ortholog Por1, members of a large superfamily of beta barrel proteins (Zeth, 2010; Zeth & Zachariae, 2018), as phospholipid scramblases. Our results indicate, that on specific dimerization, these proteins play a prominent role in transporting anionic and zwitterionic phospholipids across the OMM (Fig. 1A), which is required for mitochondrial membrane biogenesis and mitochondrial function. The unitary rate of VDAC-dimer-mediated scrambling (>10^3^ s^-1^ in reconstituted vesicles) and the abundance of VDACs (>20,000 copies per yeast cell), ensure that VDACs provide the OMM with orders of magnitude more scramblase activity than is needed to supply the mitochondrial network (0.25 fmol phospholipids per yeast cell) with lipids for cell growth and homeostasis (Petrungaro & Kornmann, 2019). As all phospholipid scramblases identified so far are α-helical membrane proteins, our results introduce the transmembrane β-barrel as a new scramblase-active protein fold. The current mechanistic paradigm for α-helical scramblases is that they provide a polar transbilayer pathway which allows phospholipids to traverse the membrane with their headgroup residing within the pathway while their acyl chains extend into the hydrophobic membrane core, as suggested by the credit-card model (Pomorski & Menon, 2006). This polar pathway can be formed by two or three adjacent membrane helices within a protein monomer as seen for GPCR and TMEM16 scramblases (Kalienkova *et al*, 2021; Khelashvili & Menon, 2022), between protomers in an oligomer as in bacteriorhodopsin and Atg9 (Maeda *et al*, 2020; Matoba *et al*, 2020; Verchere *et al*, 2017), or at the surface of the protein (Kalienkova *et al*., 2021; Maeda *et al*., 2020; Malvezzi *et al*, 2018). A polar pathway on the surface of VDAC1 and VDAC2, centered on the membrane facing glutamate, clearly lowers the barrier for lipid scrambling (Fig. S1) but is not sufficient to support fast scrambling (Fig. S6). In contrast, the dimer-1 interface depresses the free energy barrier far more substantially, enabling fast lipid scrambling. The effectiveness of the dimer-1 interface in promoting fast scrambling derives from the synergistic effect of attractive interactions between lipid headgroups and protein residues, and barrier lowering caused by membrane thinning and associated water defects (Fig. S10)(Bennett & Tieleman, 2014). Neither feature alone is sufficient to account for fast scrambling activity. Notably, because of the requirement for quaternary structure in generating a fast scrambling path, it is possible that scrambling at the OMM in cells may be regulated through control of dimerization by other proteins as well as by the lipid environment (Betaneli *et al*., 2012; Chadda *et al*, 2021; Shoshan-Barmatz *et al*., 2013).

Elimination of both VDAC homologs in yeast slowed scrambling across the OMM by more than an order of magnitude (Fig. 5D), indicating that these proteins are the principal lipid importers of the OMM. Consistent with this, depletion of VDACs in yeast and mammalian cells caused a reduction in the levels of cardiolipin (Broeskamp *et al*, 2021; Miyata *et al*, 2018), a lipid whose synthesis requires the import of phospholipid precursors from the ER. However, the residual (<10% of total) scramblase activity in *por111por211* mitochondria (Fig. 5D) suggests that there are additional scramblases. Possible candidates are the ubiquitous OMM translocation channel Tom40, a 19-stranded β-barrel which is evolutionarily related to VDAC and shares the same fold (Araiso *et al*, 2019; Tucker & Park, 2019; Wang *et al*, 2020; Zeth, 2010). In support of this idea, Tom40 dimers -like VDAC dimers-have been shown to influence the membrane through bending and destabilization (Araiso *et al*., 2019; Tucker & Park, 2019; Wang *et al*., 2020; Zeth, 2010). Other transmembrane β-barrels of the OMM like Sam50 or Mdm10 might show similar properties (Egea, 2021; Takeda *et al*, 2021). The OMM protein MTCH2, recently characterized as a protein insertase involved in the insertion of a variety of α-helical proteins into the OMM (Guna *et al*, 2022), has a prominent transmembrane groove containing polar/charged residues, reminiscent of the groove implicated in scrambling lipids according to the credit card model (Pomorski & Menon, 2006). Our CGMD analyses of monomeric MTCH2 indicate that it scrambles phospholipids at a rate similar to that of dimer-1 (Fig. 4C)(LB, AKM, RV, unpublished). Further work will be needed to quantify the relative contribution of MTCH2 to scrambling across the OMM. Recent discoveries have advanced the concept that lipid transfer proteins directly engage scramblases (Adlakha *et al*, 2022; Ghanbarpour *et al*, 2021). Thus, Vps13 and the ERMES complex that are involved in lipid flow between the ER and mitochondria physically interact with the yeast OMM proteins Mcp1 and Mdm10, respectively (Adlakha *et al*., 2022). Purified Mcp1 was recently shown to have scramblase activity (Adlakha *et al*., 2022) and it remains to be seen whether this is also the case for Mdm10. In agreement with this concept, previous studies placed VDAC at ER-mitochondria contact sites and at hubs of lipid synthesis and distribution (Broeskamp *et al*., 2021; Miyata *et al*., 2018; Tamura *et al*., 2020). Thus, in yeast, the VDAC homolog Por1 is thought to interact with the ERMES complex, the Ups-Mdm35 lipid transfer complex as well as Mdm31/32, the latter two being important for phospholipid transport across the IMS (Broeskamp *et al*., 2021; Miyata *et al*., 2018; Tamura *et al*., 2020). In this scenario, VDAC constitutes a nexus between the phospholipid transport machineries on both sides of the OMM, scrambling phospholipids to enable their entry(exit) into(from) these machineries.

## MATERIALS AND METHODS

### VDAC purification

VDAC proteins were expressed and purified as described by Dadsena *et al* (2019). Chemically competent *E. coli* BL21(DE3)-omp9 cells [F^-^, ompT hsdS_B_ (r_B-_ m_B-_) gal dcm (DE3) ΔlamB ompF::Tn5 ΔompA ΔompC ΔompN::Ω] after Prilipov *et al* (1998) were transformed with pCold vectors encoding N-terminal hexahistidine tagged human VDAC1 and VDAC2 isoforms, the point mutant VDAC2 E84L, or the pentamutant VDAC1-5V (T60V, Y62V, E73V, T83V, S101V). All constructs were verified by Sanger sequencing. For VDAC purification, the transformed cells were grown at 37°C to OD_600_ of 0.6 in 2xYT medium (16 g/l Tryptone, 5 g/l NaCl, 10 g/l Yeast extract) containing 100 µg/ml ampicillin. The cultures were cooled before adding 1 mM IPTG and incubating overnight at 15°C. Cells were harvested by centrifugation and the cell pellet was resuspended in TEN-buffer (50 mM Tris-HCl pH 8.0, 100 mM NaCl) containing a protease inhibitor cocktail (30ml of resuspension for 2 L starting cell culture). The cells were disrupted using a probe sonicator and inclusion bodies were collected by centrifugation at 25,000 x g for 30 min at 4°C. Residual cell membranes were removed by washing the pellet thrice with 15 ml TEN-buffer containing 2.5% v/v Triton X-100, using a Teflon pestle for resuspension followed by centrifugation at 25,000 x g 15min 4°C, after which the Triton X-100 was removed by further washing the inclusion bodies thrice in TEN-buffer. Washed inclusion bodies were resuspended in 4 ml TEN buffer for 2 L starting cell culture and 2 ml were used for immediate denaturing by dropwise addition into denaturation buffer (25 mM Na^+^PO_4_ pH 7.0, 100 mM NaCl, 6 M guanidine hydrochloride, 1 mM EDTA, 10 mM DTT) to a 10-times dilution. After overnight incubation at 4°C under constant stirring, the proteins were refolded by serial 10-times dilutions into first 25 mM Na^+^PO_4_ pH 7.0, 100 mM NaCl, 1 mM EDTA, 2.22% LDAO followed by overnight incubation. This material was further 10-times diluted into 25 mM Na^+^PO_4_ pH 7.0, 10 mM NaCl, 1 mM EDTA, 0.1% LDAO. After incubation at 4°C for 4h or overnight, the solution was filtered, applied onto a cation-exchange column (HiTrap^TM^ SP HP 5ml (GE Healthcare)), and eluted with a salt gradient (10 mM to 1 M NaCl in the buffer 25 mM Na^+^PO_4_ pH 7.0, 1 mM EDTA, 0.1% LDAO, 1 mM DTT). Protein-containing fractions were pooled, concentrated to 500 µl using Amicon Ultra-4 (Millipore) centrifugal filters, and loaded onto a Superdex 200 10/300 25ml size-exclusion column using SEC buffer (10 mM Tris-HCl pH 8.0, 100 mM NaCl, 0.05% LDAO). Peak fractions were pooled, quantified by absorbance (VDAC1 ɛ_280_= 38,515 M^-1^.cm^-1^, VDAC2 ɛ_280_= 37,400 M^-1^.cm^-1^) and BCA colorimetric assay, assessed for purity by Coomassie-stained SDS-PAGE and analyzed by Circular Dichroism spectroscopy using Aviv 410 CD instrument. For storage at −80°C, 10% glycerol was added, and the protein was snap frozen. The typical protein concentration obtained was 1 mg/ml from 1 L of starting culture.

### Yeast VDAC (Por1) purification

*S. cerevisiae* W303 cells were transformed with a pBEVY-GL vector (Miller et al, 1998) for galactose-inducible expression of C-terminal Twin-Strep-tagged yeast VDAC (Por1-TS), with a TEV protease cleavage site in the linker region between the Por1 sequence and the Twin-Strep tag. The expression construct was a kind gift from Dr. Susan Buchanan (National Institute of Diabetes and Digestive and Kidney Diseases). An overnight culture of the cells in synthetic complete medium minus leucine (SC-Leu) with 2% (w/v) glucose was used to inoculate 500 ml SC-Leu with 3% (w/v) glycerol and 0.1% (w/v) glucose. After overnight growth, cells were pelleted and washed twice in a small volume of yeast extract-peptone (YP) medium (1% w/v yeast extract, 2% w/v peptone) with 2% (w/v) galactose to remove all remaining glucose from the growth medium. Resuspended cells were transferred to 1.5 L of YP galactose (starting OD_600_ ∼0.6) and grown overnight at 25°C (20 hrs).

Cells were harvested, resuspended to a final volume of 50 ml in 50 mM HEPES pH 7.5, 150 mM NaCl, 1mM EDTA, 0.1 mM PMSF, Roche EDTA-free Mini cOmplete PIC tablets, and lysed with 5 passes through an Emulsiflex-C3 homogenizer at 25,000 psi. Debris and unbroken cells were removed by centrifuging for 10 min at 3000 x g, and the supernatant was subsequently centrifuged at 40,000 rpm for 1.5 h in a Beckman 45 Ti rotor to pellet membranes. The pellet was resuspended in 50 ml 50 mM HEPES pH 7.5, 150 mM NaCl, 1 mM EDTA and homogenized 10 ml at a time in a Dounce homogenizer. LDAO (1% (w/v)) was added, and the sample was stirred for 1.5 h at 4°C. Insoluble material was pelleted at 30,600 rpm for 30 minutes in a 45 Ti rotor. The supernatant was vacuum filtered through a 0.22 µm membrane and subjected to Twin-Strep-tag/Strep-Tactin XT affinity purification as follows. A 1 ml Strep-Tactin XT 4Flow high-capacity cartridge (IBA Lifesciences 2-5027-001) was washed with 14 column volumes of wash buffer (50 mM Hepes pH 7.6, 150 mM NaCl, 1 mM EDTA, 0.1% LDAO) on an ӒKTA pure™ chromatography system. The 50 ml filtered sample was applied to the resin at 1 ml/min, and then the resin was washed with wash buffer until the absorbance at 280 nm had stabilized. Flow through and wash buffer were collected for SDS-PAGE analysis. Bound Por1-TS was eluted with the addition of 50 mM Biotin in wash buffer (pH 7.6) at 1 ml/min, collecting 1 ml fractions, until absorbance at 280 nm reached baseline. Fractions were analyzed by Coomassie-stained SDS-PAGE. Peak fractions were pooled and concentrated to 500 μl using Amicon Ultra-15 10K MWCO centrifugal filters (Merck Millipore UFC901008; final LDAO = 2%) and further purified by size-exclusion chromatography as described for human VDAC constructs but with 20 mM HEPES pH 7.6, 150 mM NaCl, 0.05% LDAO as SEC buffer. Por1-TS-containing SEC fractions were concentrated to 200 ng/μl (quantified by comparison with BSA standards on Coomassie-stained SDS-PAGE) and snap-frozen in 10% (w/v) glycerol for storage at −80°C. Mass spectrometric analyses of the purified sample were performed at the Weill Cornell Medicine Proteomics and Metabolomics Core Facility.

### Pet464 purification

The purified Pet464 protein in 50 mM Tris, pH 8.0, 150 mM NaCl, 0.05% LDAO, and the corresponding plasmid pET22bbPet464ββ containing the Pet protein with a passenger domain truncation to 464 amino acids was a kind gift from Matthew Johnson and Denisse Leyton of the Australian National University (Yuan *et al*., 2018). The protein was prepared as described (Yuan *et al*., 2018). Formation of inclusion bodies was induced in BL21(DE3) cells using 0.5 mM IPTG for 4h. Cells were resuspended in 50 mM Tris-HCl pH 8.0, 150 mM NaCl, 1% Triton X-100 and pretreated with 0.7 mg/ml lysozyme chloride on ice for 10 minutes, and another 30 minutes after the addition of DNase1 and 5 mM MgCl. After cell lysis by tip sonication, inclusion bodies were collected by centrifugation at 10,000 x g, 10 min and washed three times in 50 mM Tris-HCl pH 8.0, 150 mM NaCl, 1% Triton X-100 with a final wash excluding Triton X-100. The inclusion bodies were solubilized in 50 mM Tris-HCl, pH 8.0, 8 M urea for 2 h on a rotation wheel and subsequentially centrifuged at 30,000 x g, 30 min to remove aggregates. The unfolded protein was snap frozen, and stored at −80°C. The protein was refolded by a rapid 10-fold dilution into pre-warmed refolding buffer (50 mM Tris-HCl pH 8.0, 150 mM NaCl, 0.5% LDAO) at 35°C for 10 min under constant shaking. Finally, the β-barrel was purified using first Qiagen Ni-NTA Resin Beads and further size exclusion chromatography in SEC-buffer (50 mM Tris-HCl pH 8.0, 150 mM NaCl, 0.05% LDAO). Fractions were collected, quantified by a wavelength scan (ɛ280= 48,360 M^-1^ cm^-1^), SDS-PAGE, and a BCA protein determination before further use for protein reconstitution.

### Large unilamellar vesicles (LUVs)

POPC and POPG (1435 µl and 160 µl of 25 mg/ml stock solutions in chloroform), together with (1-myristoyl-2-C_6_-NBD-PC) (195 µl of a 1 mg/ml stock solution in chloroform) as indicated, were added to a round-bottom flask. The solvent was evaporated using a rotary evaporator and the flask was placed overnight in a desiccator attached to house vacuum. The dried lipid film was resuspended by swirling in 10 ml reconstitution buffer (10 mM MOPS/Tris pH 7.0, 100 mM KCl) or crosslinking buffer (10 mM MOPS/KOH pH 7.0, 100 mM KCl) as indicated (lipid concentration 5.25 mM), and incubated at 37°C for 20 min. The resuspended lipids were briefly sonicated in a bath sonicator, before being sequentially extruded through 0.4 µm and 0.2 µm membranes in a high-pressure lipid extruder (Northern Lipids). The size (150 nm) and polydispersity index (12.5) of the preparation were determined by Dynamic Light Scattering (DLS) with the Litesizer^TM^ 500 Instrument. The liposomes had a typical concentration of 3 mM.

### VDAC reconstitution into LUVs

VDAC was reconstituted into pre-formed LUVs by a modification of the method of Brunner & Schenck (2019). Briefly, 800 µl LUVs were destabilized by adding 16 µl 10% w/v Triton X-100 and incubating for 20 min with agitation. VDAC samples were supplemented to 1.05% LDAO and agitated (600 rpm) on a thermomixer at room temperature or, in some instances at 37°C for VDAC1. The desired concentration of protein in a maximal volume of 100 µl was added to the destabilized vesicles, and the volume made up to 1 ml with SEC buffer or crosslinking-buffer (10 mM MOPS/KOH pH 7.0, 100 mM KCl) containing 0.05% LDAO as indicated, with additional LDAO to ensure equal detergent concentrations in all samples. Samples were incubated for 1 h with end-over-end mixing at room temperature (VDAC2) or in some instances agitated (600 rpm) at 37°C in the case of VDAC1. Washed Bio-Beads (140 mg) were added, and the samples were agitated (600 rpm) for 20 min at 37°C in the case of VDAC1, then transferred to the cold room for overnight incubation at 4°C with end-over-end mixing. Reconstituted vesicles were separated from Bio-Beads and used immediately for further assays. Protein concentration was determined by SDS-PAGE (Coomassie or Fluorescence Protein gel stain (Lamda Biotech)) in comparison with standards. Lipid concentration was determined by colorimetric assay of lipid phosphorus after the protocol of Rouser *et al* (1970). The protein/phospholipid ratio (PPR, mg/mmol) of the samples was calculated using experimentally determined values. PPR* (Ploier *et al*., 2016; Verchere *et al*., 2017) describing the protein per phospholipid ratio normalized against the fraction of vesicles that cannot be populated by proteins was calculated as 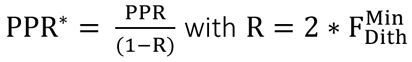 as clarified in the following section.

### Fluorescence assays for channel and scramblase activity

NBD-PC-containing proteoliposomes were assayed for VDAC channel activity with the dithionite assay after the protocol of Menon *et al*. (2011) and scramblase activity with the BSA-back extraction assay after the protocol of Kubelt *et al*. (2002). Proteoliposomes were diluted 40-times into HEPES buffer (50 mM HEPES pH 7.4, 100 mM NaCl) in a fluorimeter cuvette, and fluorescence was monitored under constant stirring (900 rpm) at 20°C in a temperature-controlled Spectrofluorometer FluoroMax + instrument using l_ex_=470 nm, l_em_=530 nm, slit widths 2.5 nm. The sample was equilibrated for 5-10 min before proceeding with the assays. For the dithionite assay, 40 µl of 1 M sodium dithionite, freshly prepared in unbuffered 0.5 M Tris, was added. For the BSA-back extraction assay, 40 µl 75 mg/ml fatty acid-free BSA (Calbiochem) in HEPES buffer was added. Collected fluorescence traces were normalized to the initial value F_max_. The fraction (f) of vesicles containing either channel or scramblase activity was calculated as 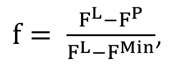 with the normalized fluorescence (F) at 350 sec after dithionite (f_Pore_) or BSA (f_Scr_) addition for protein-free (F^L^) or protein-containing liposomes (F^P^) and F^min^ as the lowest fluorescence signal detectable if all vesicles possess activity while accounting for refractory liposomes. For the dithionite assay, 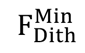 was experimentally determined using the average fluorescence value of proteoliposomes reconstituted at a protein:phospholipid ratio 1:6,000, using 15 µg/ml of protein to obtain a theoretical copy number of 30 proteins per vesicle. If the dithionite assay captures all liposomes containing VDAC proteins, the fluorescence signal describing 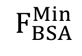 for all proteoliposomes theoretically capable of scramblase activity can be calculated as:

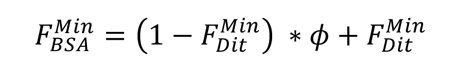

where ϕ is the fluorescence of NBD-PC when it is complexed with BSA compared with its value in the membrane. The value of ϕ was experimentally determined to be 0.4 using the fluorescence signal after 350 sec for protein-free liposomes and comparing dithionite and BSA-back extraction traces with rearrangement of the above equation as follows:

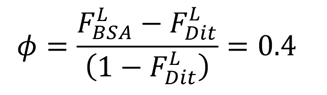

The fraction of VDAC proteoliposomes that possess scramblase activity (Q) was determined as 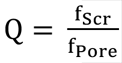. Additionally, fluorescence traces were analyzed using one-phase or two-phase exponential decay functions as determined using F-test with GraphPad Prism.

To determine oligomeric species that are reconstituted per vesicle, f_Pore_ was plotted against PPR*, and analyzed according to the Poisson model (Goren *et al*., 2014; Menon *et al*., 2011; Ploier *et al*., 2016) by fitting to a one-phase exponential function. The fit constant τ (in units of µg protein/µmol lipids) corresponds to the reconstitution condition where each vesicle has a single functional unit on average. Thus, the protein copy (C) number per 1 µmol lipids was determined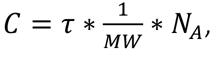 with MW = 31 kDa, and the number of vesicles (V) in 1 µmol was determined as 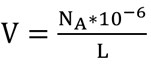 with lipids per vesicle (L) as 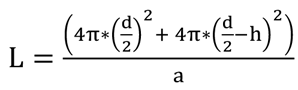 with diameter (d) as 150 nm, bilayer thickness (h) as 5 nm and phospholipid headgroup area (a) as 0.71 nm^2^ (Mimms *et al*, 1981). Proteins incorporated per vesicles are thus,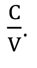.

### Scramblase assay using PI-PLC

VDAC proteoliposomes were prepared as described above except that 20,000 cpm per ml of [^3^H]inositol-labeled phosphatidylinositol ([^3^H]PI) (American Radiolabeled Chemicals) was included during the reconstitution step. [^3^H]PI was dried under nitrogen stream and taken up in 50 µl per 1 ml sample preparation in reconstitution buffer (10 mM MOPS/Tris pH 7.0, 100 mM KCl) containing 0.2% w/v Triton X-100. No NBD-PC was present in the samples. Scramblase activity was assayed using PI-specific phospholipase C (PI-PLC purchased from Sigma) as described by Wang *et al*. (2018). To 100 µl aliquots of proteoliposomes 10 µl HEPES buffer (50 mM HEPES pH 7.4, 100 mM NaCl) (or 10 µl 10% w/v Triton X-100 for sample disruption) was added. Finally, 3 µl PI-PLC working solution (10-times dilution into HEPES buffer) was added and the samples were incubated for the indicated time at 25°C. The reactions were stopped by the addition of ice-cold trichloroacetic acid to a final concentration of 12% (w/v) and placement on ice. Cytochrome c (Sigma) was added to a final concentration of 150 µg/ml for better visualization of the protein pellet. Samples were incubated on ice for 30 min before being microfuged to pellet precipitated material (including non-hydrolyzed [^3^H]PI). The supernatant containing released [^3^H]inositol-cyclic phosphate was taken for liquid scintillation counting. Control samples included proteoliposomes disrupted with Triton X-100, proteoliposomes not treated with PI-PLC and protein-free liposomes. Scintillation counts were offset-corrected using the value obtained from non-treated samples, and data were subsequently normalized to the maximum extent of hydrolysis determined from Triton X-100 disrupted samples. The percentage of hydrolyzed PI was graphed as a function of time and traces were fitted to a one-phase exponential association using GraphPad Prism software.

### EGS crosslinking

Ethylene glycol bis(succinimidyl succinate) (EGS; ThermoFisher) crosslinking was done according to the protocol of Zalk *et al*. (2005) and Kim *et al*. (2019) with modifications. VDAC was crosslinked in LDAO or after reconstitution into LUVs prepared in crosslinking buffer (10 mM MOPS/KOH pH 7.0, 100 mM KCl). Samples to be crosslinked in LDAO were applied to a desalting spin column for buffer exchange to crosslinking buffer containing 0.05% LDAO (additional LDAO was added as indicated). EGS was dissolved in DMSO, diluted 20-times in crosslinking buffer (with additional LDAO as indicated), before being diluted 10-times into VDAC samples (15 µg/ml or 150 µg/ml when used for further reconstitution) to achieve a final 16x mole excess over reactive sites (lysine residues + N-terminus), with final DMSO = 0.5%. Non-crosslinked controls were treated with an equivalent amount of DMSO. After agitation (1000 rpm) in a thermomixer at 20°C for 40 min, the reaction was stopped by adding 5xLaemmli buffer to 1x (60 mM Tris-HCl pH 6.8, 2% w/v SDS, 10% v/v glycerol, 5% v/v β-mercaptoethanol, 0.01% bromophenol blue) and heating to 95°C for 3 min. Samples were taken for immunoblotting using anti His-tag antibody (ThermoFisher). Crosslinking and control reactions intended for scramblase activity assays were stopped with 1 M Tris pH 7 to a final concentration of 10 mM, instead of Laemmli buffer, and used for reconstitution into LUVs whilst ensuring the same DMSO and EGS concentrations in control protein-free liposomes. The same protocol was followed for crosslinking with dithiobis(succinimidyl propionate) (DSP; ThermoFisher) and samples were taken up in 4xSDS-PAGE buffer to 1x (75 mM Tris-HCl pH 6.8, 2% w/v SDS, 10% glycerol, 0.05% bromophenol blue) instead of Laemmli buffer for immunoblotting. For cleavage of DSP, 50 mM DTT (from 1 M stock prepared in reconstitution buffer) was added and the samples were incubated for 30 min at 37°C while agitated (600 rpm). Control and non-cleaved samples were treated equally omitting DTT.

### Assay of slow scrambling

To assay slow scrambling, proteoliposomes were prepared without NBD-PC and were subsequently asymmetrically labeled by adding NBD-PC (from an EtOH stock solution) to a final concentration of 0.25 mol% lipid, and final EtOH concentration of 0.4 %. After 30 min incubation on ice, the samples were shifted to 20°C and 50 µl aliquots were taken for BSA-back extraction assay as described above every 2 h for up to 30 h. Normalized fluorescence signal 100 sec after BSA addition was used to determine the fraction of NBD-analogs at the outer leaflet setting time point 0 as 100% of analogs present at the outer leaflet. Traces were fitted to a one-phase exponential decay function using GraphPad Prism software.

### Isolation of yeast mitochondria

Crude mitochondria were prepared from yeast after the protocol of Daum *et al* (1981) with some modifications. Briefly, yeast cells were pre-grown in YPD (1% w/v yeast extract, 2% w/v peptone, 2% w/v dextrose) over a weekend, 10-times diluted into rich-lactate (RL) media (1% w/v yeast extract, 2% w/v peptone, 0.05% w/v dextrose, 2% v/v lactic acid, 3.4 mM CaCl2 2H_2_0, 8.5 mM NaCl, 2.95 mM MgCl_2_ 6H_2_0, 7.35 mM KH_2_PO_4_, 18.7 mM NH_4_Cl, pH 5.5 adjusted with KOH) containing 2 mM ethanolamine and grown for 4 h at 30°C. 6 OD_600_ were added to 500 ml RL containing 2 mM ethanolamine and further grown overnight to an OD_600_ of 4 to 5. Cells were harvested, washed with 1 mM EDTA and taken up in T-buffer (0.1 M Tris-HCl, 10 mM DTT, pH 9.4) at 0.5 g cell pellet/ml. Cells were incubated for 10 min at 30°C, harvested and washed in 30°C warm S-buffer (1.2 M sorbitol, 20 mM KH_2_PO_4_, pH 7.4 with KOH) at 0.15 g cell pellet/ml. Cells were collected and taken up in S-buffer containing 1 mg Zymolyase per g yeast cell pellet at 0.3 g cell pellet/ml and incubated at 30°C for 1 h. Spheroplasts were diluted in equal volume of ice-cold S-buffer and harvested at 4°C. Cells are resuspended in ice-cold D_-_-buffer (10 mM MES, 0.6 M sorbitol, pH 6 with NaOH) containing 0.2% w/v fatty acid free BSA and 1 mM PMSF at 0.75 g cell pellet/ml, and disrupted with 10 strokes of a Dounce homogenizer. Mitochondria were collected by differential centrifugation, first, intact cells were removed by two slow spins at 1,400 x g 4°C for 5 min. Crude mitochondria were pelleted by centrifugation at 10,000 x g 12 min 4°C. Mitochondria were taken up in D_-_-buffer, snap frozen and stored at −80°C. The protein concentration of the preparations was determined using BCA protein assay in the presence of 0.8% SDS and Coomassie staining.

### Transport-coupled PS decarboxylation by yeast mitochondria

Conversion of NBD-PS to NBD-PE by crude mitochondria was assayed according to the protocol of Miyata *et al* (2016) with modifications. Crude mitochondria were diluted to 5 mg/ml in D_-_-buffer. The assay was started by 5-times dilution of the mitochondria into D_+_-buffer (10 mM MES, 0.6 M sorbitol, pH 6 with NaOH, 12.5 mM EDTA) containing 1.25 µM 16:0 – 6:0 NBD-PS (Avanti) and 0.3 µM C6-NBD-PC. Samples were incubated at 30°C and 100 µl aliquots were taken at the respective time points. To stop the reaction, 750 µl CHCl3:MeOH v:v 1:2 was added. Subsequentially, 250 µl CHCl_3_ and 250 µl 0.2 M KCl were added, and the samples were centrifuged at 2,200 x g for 5 min at RT. The CHCl_3_ phase was collected and dried under nitrogen. Samples were taken up in MeOH and spotted onto an activated thin layer chromatography (TLC) silica gel 60 plate (Merck). The plate was developed in CHCl_3_:MeOH:Acetic Acid:H_2_0 (25:15:4:2, by volume) and NBD fluorescence was visualized using the BioRad ChemiDoc System. Densitometry of the fluorescence (F) signal was used to determine PS to PE conversion according to the equation 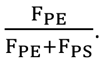 Psd1 and Tom70 levels were determined by immunoblot to establish similar content of these OMM and IMM markers in the assay samples.

### Analysis of the kinetics of transport-coupled PS decarboxylation by yeast mitochondria

The kinetic model is described in Fig. 5A - the model specifies 4 pools of lipid, and 5 rate constants. We took advantage of the environment sensitivity of NBD-PS fluorescence (∼16-fold higher fluorescence of NBD-PS in a membrane environment compared with that of an NBD-PS monomer in solution (Fig. S12G)) to estimate the effective OMM adsorption and desorption rate constants (k_0_ and k_1_). Upon mixing NBD-PS with an excess of liposomes, fluorescence increased exponentially (Fig. S12H) with a rate set by the sum of both k_0_ and k_1_. The desorption rate constant k_1_ was separately determined using BSA-back extraction with asymmetrically NBD-PS-labeled liposomes and excess BSA such that k_1_ was rate-limiting, producing a single exponential time course (Fig. S12I)(Panagabko *et al*, 2019). Taken together, these measurements provided estimates of k_0_ ∼ 1 sec^-1^ and k_1_ ∼ 0.05 sec^-1^. As NBD-PS was added to mitochondria above its critical micelle concentration (CMC) of 0.165 µM (Wustner *et al*, 2000; Yegneswaran *et al*, 2003), aqueous NBD-PS will equilibrate between monomeric [PS] and micellar [M] states. Only monomeric NBD-PS can be directly incorporated into lipid bilayers, and therefore the effective rate constant k_0_ in our kinetic model represents a simplified composite of both the desorption of NBD-PS monomers from a micellar pool as well as their subsequent incorporation into membranes. Although the desorption rate constant k_1_ may have been overestimated due to contributions by a micellar pool of NBD-PS driving the BSA-back extraction together with preloaded liposomes, this is unlikely as similar values of k_1_ were measured when using NBD-PS concentrations both above and below the CMC, indicating that no residual micelles were present after NBD-PS incorporation into liposomes (Fig. S12J). For the lipid scramblase step, rate constants k_2_ and k_3_ were assumed to be equal as lipid scrambling is a passive bidirectional transport step. We also assumed that these rate constants are larger than 0.01 sec^-1^ in wild-type mitochondria based on previous studies (Janssen *et al*., 1999). Finally, the effective rate constant k_4_ was left unconstrained for fits to the PE production time course using wild-type mitochondria and estimated together with k_2_ and k_3_ via a systematic parameter search that minimized the error between the model and experimental time courses using custom-written routines in Igor Pro 8 (Wavemetrics). The four-state kinetic model was implemented using the Bulirsch-Stoer method with Richardson extrapolation for numerical integration.

The best-fitting model for the data obtained with wild-type mitochondria produced k_2_ = k_3_ = 0.027 ± 0.006 sec^-1^ and k_4_ = 0.0018 ± 0.00005 sec^-1^. As porin deletion would not be expected to alter k_0_, k_1_, or k_4_, only k_2_ and k_3_ were adjusted in the kinetic model fits for the data obtained with *por111por211* mitochondria. The best-fitting model produced k_2_ = k_3_ = 0.0025 ± 0.0004 sec^-1^ in the mutant mitochondria, indicating that the lipid scrambling rate had dropped more than 10-fold in the absence of VDAC (Fig. 5D).

### Coarse-grained molecular dynamics simulations (CGMD)

#### CGMD - General settings and system preparation

All molecular dynamics simulations were performed using the simulation package Gromacs version 5.1.4 (Abraham *et al*, 2015) with plugin Plumed version 2.3 (Tribello *et al*, 2014). We employed i) coarse-grained Martini force-field version 3.0 (Souza *et al*, 2021) and ii) coarse-grained Martini force-field version 2.2 (de Jong *et al*, 2013; Marrink *et al*, 2007; Monticelli *et al*, 2008) with ElNeDyn network (Periole *et al*, 2009). For Martini 2 simulations of VDAC dimers, the Lennard-Jones interactions between protein beads were scaled down by 10% to prevent unrealistically strong protein-protein interactions (Javanainen *et al*, 2017). We used a structure of human VDAC1 (hVDAC1) dimer solved by x-ray diffraction with a resolution of 2.7 Å and denoted as 6g6u in the RCSB Protein Data Bank (www.rcsb.org/structure/6g6u). A single monomer of VDAC1 was minimized in vacuum using atomistic Amber 99SB-ILDN force-field (Lindorff-Larsen *et al*, 2010) with steepest descent algorithm until the maximum force on any atom was lower than 100 kJ mol^-^ 1 _nm_-1_._

For Martini 3 simulations, the minimized atomistic protein structure was coarse-grained using Martinize2 script (github.com/marrink-lab/vermouth-martinize) constructing an elastic network with a force constant of 1000 kJ mol^-1^ nm^-2^ to fix the tertiary structure of VDAC1. Dimeric structures were generated by placing the coarse-grained VDAC1 monomers at appropriate positions. No restraints maintaining the dimeric structure of VDAC1 were used unless explicitly stated otherwise. Each VDAC1 structure was embedded in the center of an xy-positioned membrane using the Insane script (github.com/Tsjerk/Insane). For simulations with monomeric VDAC1, the membrane was composed of roughly 660 molecules of 1-palmitoyl-2-oleoyl-sn-glycero-3-phosphocholine (POPC), while for the simulations of dimeric interfaces, the membrane consisted of roughly 920 molecules of POPC. Systems had an approximate size of roughly 15 × 15 × 12 nm for simulations of monomeric VDAC1 and 18 × 18 × 11 nm for dimer simulations and were solvated with roughly 16,000 and 20,000 water beads, respectively. KCl ions were added to each system at a concentration of 100 mM with an excess of ions to neutralize the system. K^+^ ions were modeled as SQ5 particles with charge of +1 and mass of 39 a.u. Each constructed system was minimized using the steepest descent algorithm and a maximum force tolerance of 200 kJ mol^-1^ nm^-1^.

In Martini 2 simulations with ElNeDyn, the VDAC1 monomer and dimer structures were coarse-grained and embedded into the membrane using the CHARMM-GUI web interface (Jo *et al*, 2008). The systems with monomeric and dimeric VDAC1 had similar sizes and the same molecular composition as the Martini 3 systems. In the case of Martini 2, the solvated K^+^ ions were modeled as Qd-type particles with charge +1 (the same as Na^+^ ions). The dimeric structures of VDAC1 were not fixed by any external potentials and all Martini 2 systems were minimized the same way as Martini 3 systems.

### CGMD - Equilibration and production simulations

For both Martini 3 and Martini 2 simulations, the equilibration was performed in five stages with increasing simulation time-step (dt) and length (t): I) dt = 2 fs, t = 0.5 ns; II) dt = 5 fs, t = 1.25 ns; III) dt = 10 fs, t = 1 ns; IV) dt = 20 fs, t = 30 ns; V) dt = 20 fs, t = 15 ns. In all, but the stage V, position restraints in the form of harmonic potential with a force constant of 1000 kJ mol^-1^ nm^-2^ were applied to all coordinates of protein backbone beads. For Martini 3 systems with VDAC1 dimers, water restraints were applied in stages I-III to keep water from entering the hydrophobic core of the membrane. For this purpose, inverted flat-bottom potential with a force constant (k) of 10 kJ mol^-1^ nm^-2^ and a reference distance (ref) of 2.2 nm from the membrane center of mass was applied to the z-coordinates of all water beads. Equilibration was performed in the NPT ensemble with the temperature being maintained at 300 K using stochastic velocity-rescaling thermostat (Bussi *et al*, 2007) with a coupling constant of 1 ps. Water with ions, membrane, and protein(s) were coupled to three separate thermal baths. The pressure was kept at 1 bar using Berendsen barostat (Berendsen *et al*, 1984) with a coupling constant of 12 ps. The simulation box size was scaled semi-isotropically with a compressibility of 3 × 10^-4^ bar^-1^ in both xy-plane and z-direction.

To integrate Newton’s equations of motion, we used the leap-frog algorithm. Non-bonded interactions were cut-off at 1.1 nm. Van der Waals potential was shifted to zero at the cut-off distance. The reaction field was used to treat electrostatic interactions with a relative dielectric constant of 15. For Martini 3 simulations, LINCS (Hess *et al*, 1997) parameters lincs-order and lincs-iter were set to 8 and 2, respectively, to avoid artificial temperature gradients (Thallmair *et al*, 2021). Following equilibration, monomeric and several dimeric VDAC1 structures were simulated in Martini 3 for 10 µs using the same simulation settings as in stage V of the equilibration, except the Berendsen barostat was replaced with Parrinello-Rahman barostat (Parrinello & Rahman, 1981). The same simulation settings were also used for other simulations described in the section Free energy calculations. Three independent replica simulations were performed for each of the systems with monomeric or dimeric VDAC1. Apart from unrestrained simulations, we simulated the fastest scrambling dimer, dimer-1, restrained to its initial structure using harmonic potential (k = 1000 kJ mol-1 nm-2) in the x-and y-coordinates of backbone beads of the glutamate 73 and serine 193 of both VDAC1 protomers.

### CGMD - Analysis

For each of the three independent molecular dynamics simulations, we calculated the percentage of scrambled lipids in time, analyzing a simulation snapshot every 10 ns. Lipid was identified as “scrambled” if it was positioned in a different membrane leaflet than at the start of the simulation. Lipids were assigned to the leaflet by the position of their phosphate bead with respect to the membrane center. For further analysis, all three independent replicas were concatenated and centered on protein beads using its root-mean-square deviation. We constructed a water defect map describing membrane disruption around the protein structure. Water defect was defined as the average number of water beads located closer to the membrane center on the z axis than 1 nm. The average number was calculated using xy-mesh with 0.1 nm bin every 100 ps. The same mesh was employed to calculate the average membrane thickness using z-positions of lipid phosphates. The average membrane thickness was defined as the difference between the position of the upper-and lower-leaflet phosphates. The bins with less than 100 samples were excluded from the analysis. The code used for the analysis of lipid scrambling and membrane disruption is available from https://github.com/Ladme/scramblyzer and https://github.com/Ladme/memdian, respectively. The position of the beta-strands was approximated by the average position of the backbone bead of the central residue in the xy-plane.

### CGMD - Free energy calculations

We calculated the free energy of lipid flip-flop in several systems: a) in protein-free POPC membrane (Martini 3 & Martini 2), b) in system with VDAC1 monomer (Martini 3 & Martini 2), c) in system with VDAC1 dimer-1 (Martini 3), d) in system with VDAC1 monomer while turning off the attractive Lennard-Jones (LJ) interactions between the scrambling lipid and VDAC (Martini 3), and e) in system with VDAC1 dimer-1 while turning off the attractive Lennard-Jones interactions between the scrambling lipid and VDAC (Martini 3).

To enhance the sampling of the lipid flip-flop, we employed the umbrella sampling method (Torrie & Valleau, 1977). For free energy calculations in the presence of VDAC1 monomer, Hamiltonian replica exchange was further applied (Fukunishi *et al*, 2002). Three one-dimensional collective variables (CVs) were used to capture the lipid flip-flop in different systems. For simulations without the presence of VDAC1, the oriented z-distance between a chosen lipid phosphate bead and the local membrane center of mass was used as a CV. The local membrane center of mass was calculated from the positions of lipid beads localized inside a cylinder with a radius of 2.0 nm and its principal axis going through the selected phosphate bead. For simulations of systems with VDAC1 monomer, the CV was defined as the oriented z-distance between a chosen lipid phosphate bead and backbone bead of glutamate 73. In systems with VDAC1 dimer-1, the CV was defined as the oriented z-distance between a chosen lipid phosphate bead and the center of mass of the dimer-1 which was positioned close to the center of the membrane on the z-axis. During the pulling and umbrella sampling simulations, the dimer-1 was restrained in the same way as described in the section Equilibration and production simulations.

Initial configurations for umbrella sampling were generated by pulling the chosen lipid phosphate bead through the membrane for 1 µs with a pulling rate of 4.2 nm µs^-1^ and initial reference distance of 2.1 (for systems without VDAC1) or ±2.0 (for systems with VDAC1) nm using a harmonic potential. In all systems, one pulling simulation was run for each direction of pulling (from the upper to the lower membrane leaflet and in the opposite direction). The exception was Martini 3 systems with VDAC1 monomer, where two pulling simulations in each direction were performed to generate a larger ensemble of initial configurations for umbrella sampling with Hamiltonian replica exchange. In monomeric VDAC1 systems, the pulled phosphate was restrained by an xy-plane flat-bottom potential (k = 500 kJ mol^-1^ nm^-2^, ref = 1.5 nm) to stay close the backbone bead of the glutamate 73 residue. In dimeric VDAC1 systems, the phosphate was restrained by an xy-plane flat-bottom potential (k = 500 kJ mol^-1^ nm^-2^, ref = 2.5 nm) to stay close to the dimeric interface.

For systems with VDAC1, two independent sets of umbrella sampling windows were prepared to observe any possible hysteresis of the flip-flop process. For Martini 2 simulations, the initial configurations for each set of umbrella sampling windows were obtained from the pulling simulation performed in the specific direction and the calculation was enhanced by applying Hamiltonian replica exchange in these windows. In Martini 3 simulations with VDAC1 monomer, the efficiency of the Hamiltonian replica exchange was further enhanced by using configurations from two pulling simulations performed in the same direction. For systems without VDAC1, only one pulling direction was used.

We used between 15 (Martini 3 simulations of dimer-1) and 72 windows (Martini 2 simulations of monomeric VDAC) distributed along the range of the CV for each pulling direction. See Tables 1-5 for the complete list of umbrella sampling windows that were used for the calculations. After a short 30 ns equilibration, each window was sampled for a specific time which differed based on the system sampled and force-field used. The simulated times necessary to obtain converged free energy profiles were a) 1 µs for protein-free membrane, b) 4 or 8 µs for simulations with VDAC1 monomer in Martini 3 and Martini 2, respectively, c) 6 µs for simulations with VDAC1 dimer, d) 2 µs for simulations with VDAC1 monomer and attractive LJ interactions turned off, and e) 4 µs for simulations with VDAC1 dimer and attractive LJ interactions turned off. In simulations in which Hamiltonian replica exchange was applied, the exchange was attempted every 10,000 integration steps (200 ps). Free energy profiles were obtained from the umbrella sampling windows using the weighted histogram analysis method (Kumar *et al*, 1992; Souaille & Roux, 2001) as implemented in the Gromacs tool g_wham (Hub *et al*, 2010).

**Table 1.** Distribution of umbrella sampling windows along the collective variable with the biasing force constants used for Martini 3 and Martini 2 free energy calculations of lipid scrambling in a protein-free membrane.

| Window No. | Reference position [nm] | Force constant [kJ mol<sup>-1</sup> nm<sup>-2</sup>] | Window No. | Reference position [nm] | Force constant [kJ mol<sup>-1</sup> nm<sup>-2</sup>] |
| --- | --- | --- | --- | --- | --- |
| 1 | 2.00 | 3000 | 25 | -0.05 | 5000 |
| 2 | 1.90 | 3000 | 26 | -0.10 | 5000 |
| 3 | 1.80 | 3000 | 27 | -0.15 | 5000 |
| 4 | 1.70 | 3000 | 28 | -0.20 | 5000 |
| 5 | 1.60 | 3000 | 29 | -0.25 | 5000 |
| 6 | 1.50 | 3000 | 30 | -0.30 | 3000 |
| 7 | 1.40 | 3000 | 31 | -0.40 | 3000 |
| 8 | 1.30 | 3000 | 32 | -0.50 | 3000 |
| 9 | 1.20 | 3000 | 33 | -0.60 | 3000 |
| 10 | 1.10 | 3000 | 34 | -0.70 | 3000 |
| 11 | 1.00 | 3000 | 35 | -0.80 | 3000 |
| 12 | 0.90 | 3000 | 36 | -0.90 | 3000 |
| 13 | 0.80 | 3000 | 37 | -1.00 | 3000 |
| 14 | 0.70 | 3000 | 38 | -1.10 | 3000 |
| 15 | 0.60 | 3000 | 39 | -1.20 | 3000 |
| 16 | 0.50 | 3000 | 40 | -1.30 | 3000 |
| 17 | 0.40 | 3000 | 41 | -1.40 | 3000 |
| 18 | 0.30 | 3000 | 42 | -1.50 | 3000 |
| 19 | 0.25 | 5000 | 43 | -1.60 | 3000 |
| 20 | 0.20 | 5000 | 44 | -1.70 | 3000 |
| 21 | 0.15 | 5000 | 45 | -1.80 | 3000 |
| 22 | 0.10 | 5000 | 46 | -1.90 | 3000 |
| 23 | 0.05 | 5000 | 47 | -2.00 | 3000 |
| 24 | 0.00 | 5000 |  |  |  |

**Table 2.**
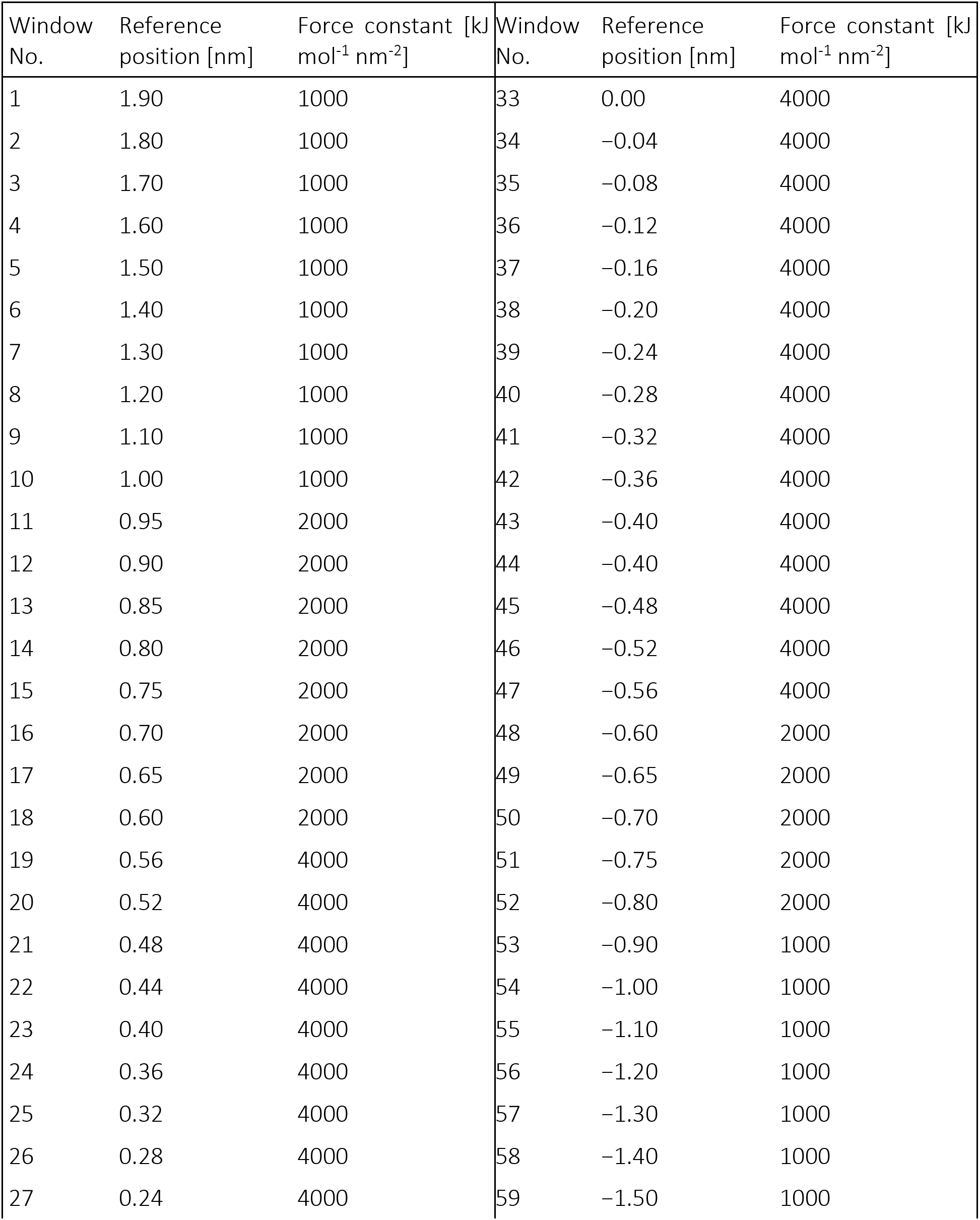
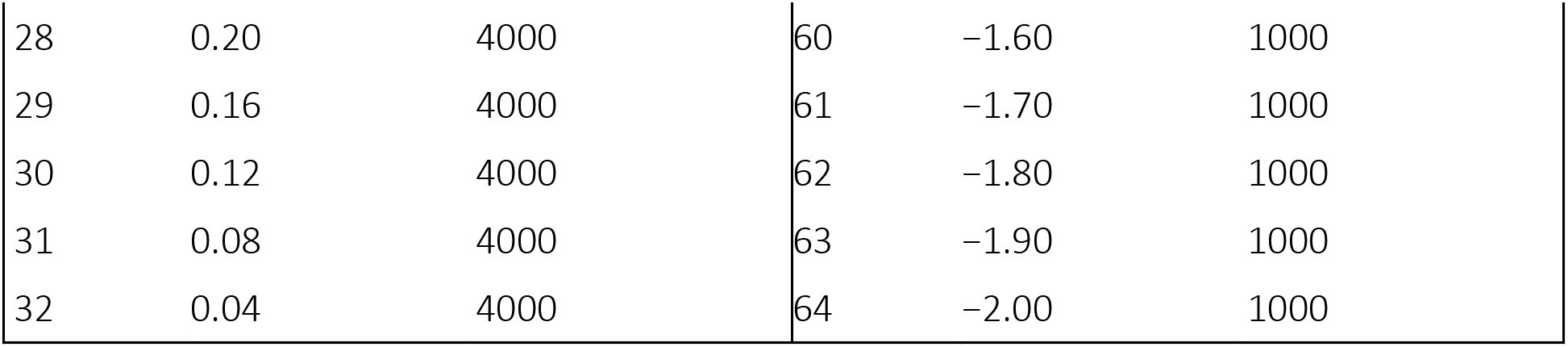
Distribution of umbrella sampling windows along the collective variable with the biasing force constants used for Martini 3 free energy calculations of lipid scrambling along monomeric VDAC1.

| Window No. | Reference position [nm] | Force constant [kJ mol<sup>-1</sup> nm<sup>-2</sup>] | Window No. | Reference position [nm] | Force constant [kJ mol<sup>-1</sup> nm<sup>-2</sup>] |
| --- | --- | --- | --- | --- | --- |
| 1 | 1.90 | 1000 | 33 | 0.00 | 4000 |
| 2 | 1.80 | 1000 | 34 | -0.04 | 4000 |
| 3 | 1.70 | 1000 | 35 | -0.08 | 4000 |
| 4 | 1.60 | 1000 | 36 | -0.12 | 4000 |
| 5 | 1.50 | 1000 | 37 | -0.16 | 4000 |
| 6 | 1.40 | 1000 | 38 | -0.20 | 4000 |
| 7 | 1.30 | 1000 | 39 | -0.24 | 4000 |
| 8 | 1.20 | 1000 | 40 | -0.28 | 4000 |
| 9 | 1.10 | 1000 | 41 | -0.32 | 4000 |
| 10 | 1.00 | 1000 | 42 | -0.36 | 4000 |
| 11 | 0.95 | 2000 | 43 | -0.40 | 4000 |
| 12 | 0.90 | 2000 | 44 | -0.40 | 4000 |
| 13 | 0.85 | 2000 | 45 | -0.48 | 4000 |
| 14 | 0.80 | 2000 | 46 | -0.52 | 4000 |
| 15 | 0.75 | 2000 | 47 | -0.56 | 4000 |
| 16 | 0.70 | 2000 | 48 | -0.60 | 2000 |
| 17 | 0.65 | 2000 | 49 | -0.65 | 2000 |
| 18 | 0.60 | 2000 | 50 | -0.70 | 2000 |
| 19 | 0.56 | 4000 | 51 | -0.75 | 2000 |
| 20 | 0.52 | 4000 | 52 | -0.80 | 2000 |
| 21 | 0.48 | 4000 | 53 | -0.90 | 1000 |
| 22 | 0.44 | 4000 | 54 | -1.00 | 1000 |
| 23 | 0.40 | 4000 | 55 | -1.10 | 1000 |
| 24 | 0.36 | 4000 | 56 | -1.20 | 1000 |
| 25 | 0.32 | 4000 | 57 | -1.30 | 1000 |
| 26 | 0.28 | 4000 | 58 | -1.40 | 1000 |
| 27 | 0.24 | 4000 | 59 | -1.50 | 1000 |
| 28 | 0.20 | 4000 | 60 | -1.60 | 1000 |
| 29 | 0.16 | 4000 | 61 | -1.70 | 1000 |
| 30 | 0.12 | 4000 | 62 | -1.80 | 1000 |
| 31 | 0.08 | 4000 | 63 | -1.90 | 1000 |
| 32 | 0.04 | 4000 | 64 | -2.00 | 1000 |

**Table 3.**
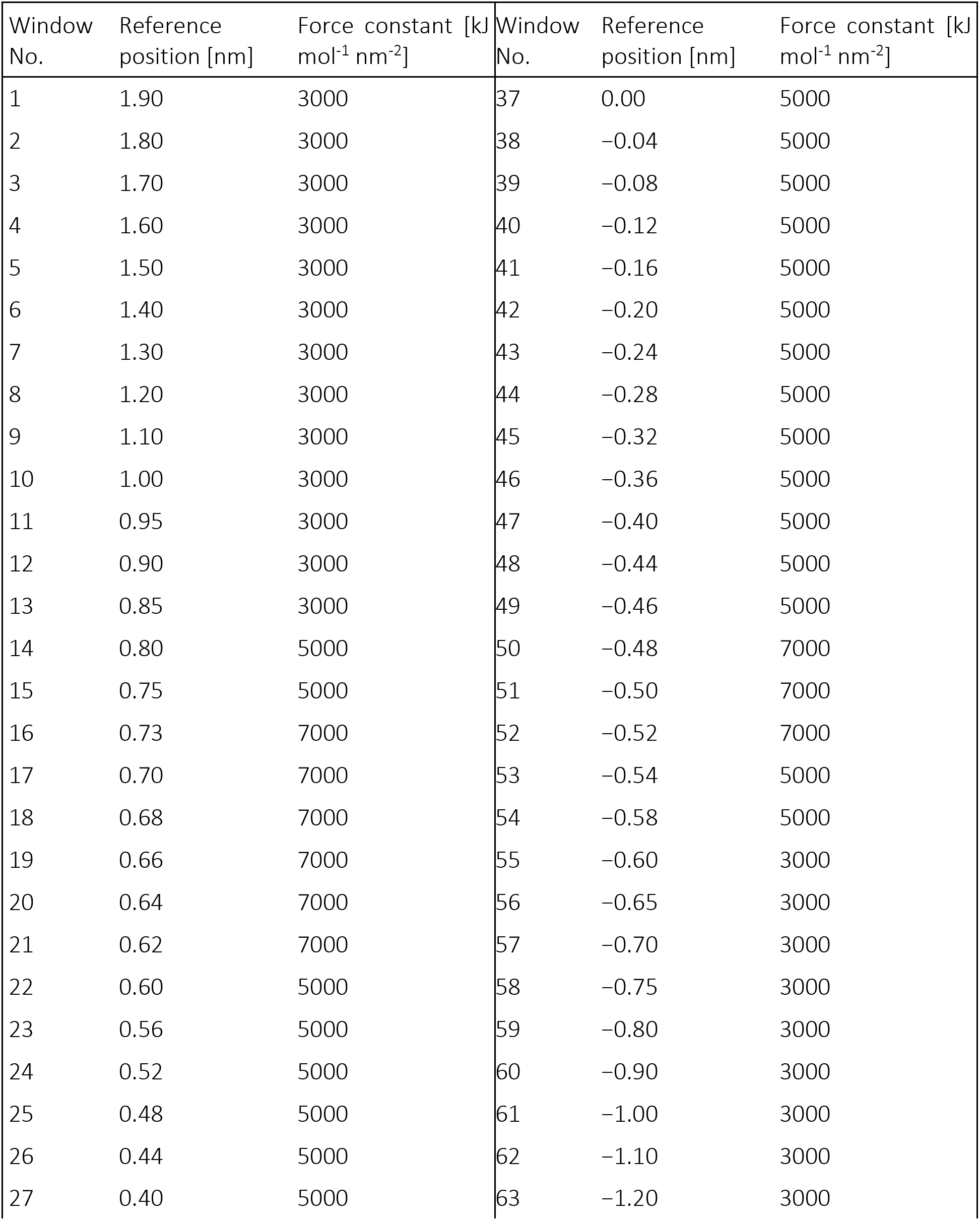
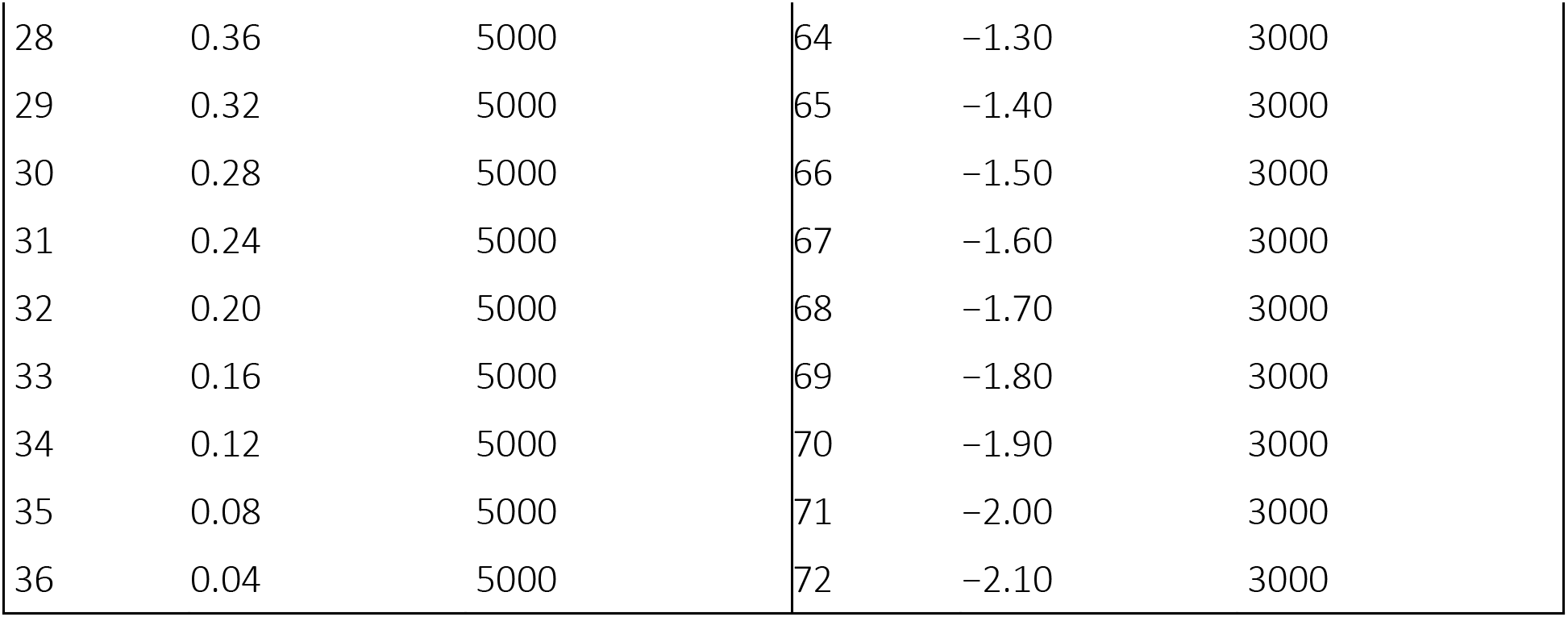
Distribution of umbrella sampling windows along the collective variable with the biasing force constants used for Martini 2 free energy calculations of lipid scrambling along monomeric VDAC1.

**Table 4.** Distribution of umbrella sampling windows along the collective variable with the biasing force constants used for Martini 3 free energy calculations of lipid scrambling along dimer-1.

| Window No. | Reference position [nm] | Force constant [kJ mol<sup>-1</sup> nm<sup>-2</sup>] | Window No. | Reference position [nm] | Force constant [kJ mol<sup>-1</sup> nm<sup>-2</sup>] |
| --- | --- | --- | --- | --- | --- |
| 1 | 1.80 | 50 | 9 | -0.20 | 200 |
| 2 | 1.40 | 50 | 10 | -0.40 | 200 |
| 3 | 1.00 | 50 | 11 | -0.60 | 100 |
| 4 | 0.80 | 100 | 12 | -0.80 | 100 |
| 5 | 0.60 | 100 | 13 | -1.00 | 50 |
| 6 | 0.40 | 200 | 14 | -1.40 | 50 |
| 7 | 0.20 | 200 | 15 | -1.80 | 50 |
| 8 | 0.00 | 200 |  |  |  |

**Table 5.** Distribution of umbrella sampling windows along the collective variable with the biasing force constants used for Martini 3 free energy calculations of lipid scrambling along VDAC1 monomer and dimer-1 with attractive LJ interactions turned off.

| Window No. | Reference position [nm] | Force constant [kJ mol<sup>-1</sup> nm<sup>-2</sup>] | Window No. | Reference position [nm] | Force constant [kJ mol<sup>-1</sup> nm<sup>-2</sup>] |
| --- | --- | --- | --- | --- | --- |
| 1 | 2.20 | 1000 | 33 | 0.04 | 4000 |
| 2 | 2.10 | 1000 | 34 | 0.00 | 4000 |
| 3 | 2.00 | 1000 | 35 | -0.04 | 4000 |
| 4 | 1.90 | 1000 | 36 | -0.08 | 4000 |
| 5 | 1.80 | 1000 | 37 | -0.12 | 4000 |
| 6 | 1.70 | 1000 | 38 | -0.16 | 4000 |
| 7 | 1.60 | 1000 | 39 | -0.20 | 4000 |
| 8 | 1.50 | 1000 | 40 | -0.24 | 4000 |
| 9 | 1.40 | 1000 | 41 | -0.28 | 4000 |
| 10 | 1.30 | 1000 | 42 | -0.32 | 4000 |
| 11 | 1.20 | 1000 | 43 | -0.36 | 4000 |
| 12 | 1.10 | 1000 | 44 | -0.40 | 4000 |
| 13 | 1.00 | 1000 | 45 | -0.44 | 4000 |
| 14 | 0.90 | 1000 | 46 | -0.48 | 4000 |
| 15 | 0.80 | 1000 | 47 | -0.52 | 4000 |
| 16 | 0.75 | 2000 | 48 | -0.56 | 4000 |
| 17 | 0.70 | 2000 | 49 | -0.60 | 2000 |
| 18 | 0.65 | 2000 | 50 | -0.65 | 2000 |
| 19 | 0.60 | 2000 | 51 | -0.70 | 1000 |
| 20 | 0.56 | 4000 | 52 | -0.80 | 1000 |
| 21 | 0.52 | 4000 | 53 | -0.90 | 1000 |
| 22 | 0.48 | 4000 | 54 | -1.00 | 1000 |
| 23 | 0.44 | 4000 | 55 | -1.10 | 1000 |
| 24 | 0.40 | 4000 | 56 | -1.20 | 1000 |
| 25 | 0.36 | 4000 | 57 | -1.30 | 1000 |
| 26 | 0.32 | 4000 | 58 | -1.40 | 1000 |
| 27 | 0.28 | 4000 | 59 | -1.50 | 1000 |
| 28 | 0.24 | 4000 | 60 | -1.60 | 1000 |
| 29 | 0.20 | 4000 | 61 | -1.70 | 1000 |
| 30 | 0.16 | 4000 | 62 | -1.80 | 1000 |
| 31 | 0.12 | 4000 | 63 | -1.90 | 1000 |
| 32 | 0.08 | 4000 | 64 | -2.00 | 1000 |

**Table 6.** Yeast strains used in this study.

| Strain name | Genetic Background | Mating Type | Genotype | Source |
| --- | --- | --- | --- | --- |
| TKY705 (Wild Type) | W303 | a | ura3-52 trp1Δ2 leu2-3_112 his3-11 ade2-1 CAN1 | Miyata et al. (2018) |
| TKY705 psd1Δ | W303 | a | TKY705, psd1Δ::natMX4 | This study |
| TKY705 por1Δ | W303 | a | TKY705, por1Δ::natMX4 | This study |
| TKY705 por2Δ | W303 | a | TKY705, por2Δ::natMX4 | This study |
| TKY705 por1Δpor2Δ | W303 | a | TKY705, por1Δ::natNT2 por2Δ::hphNT1 | Miyata et al. (2018) |

Miyata N, Fujii S, Kuge O (2018) Porin proteins have critical functions in mitochondrial phospholipid metabolism in yeast. J Biol Chem 293: 17593-17605

**Table 7.**
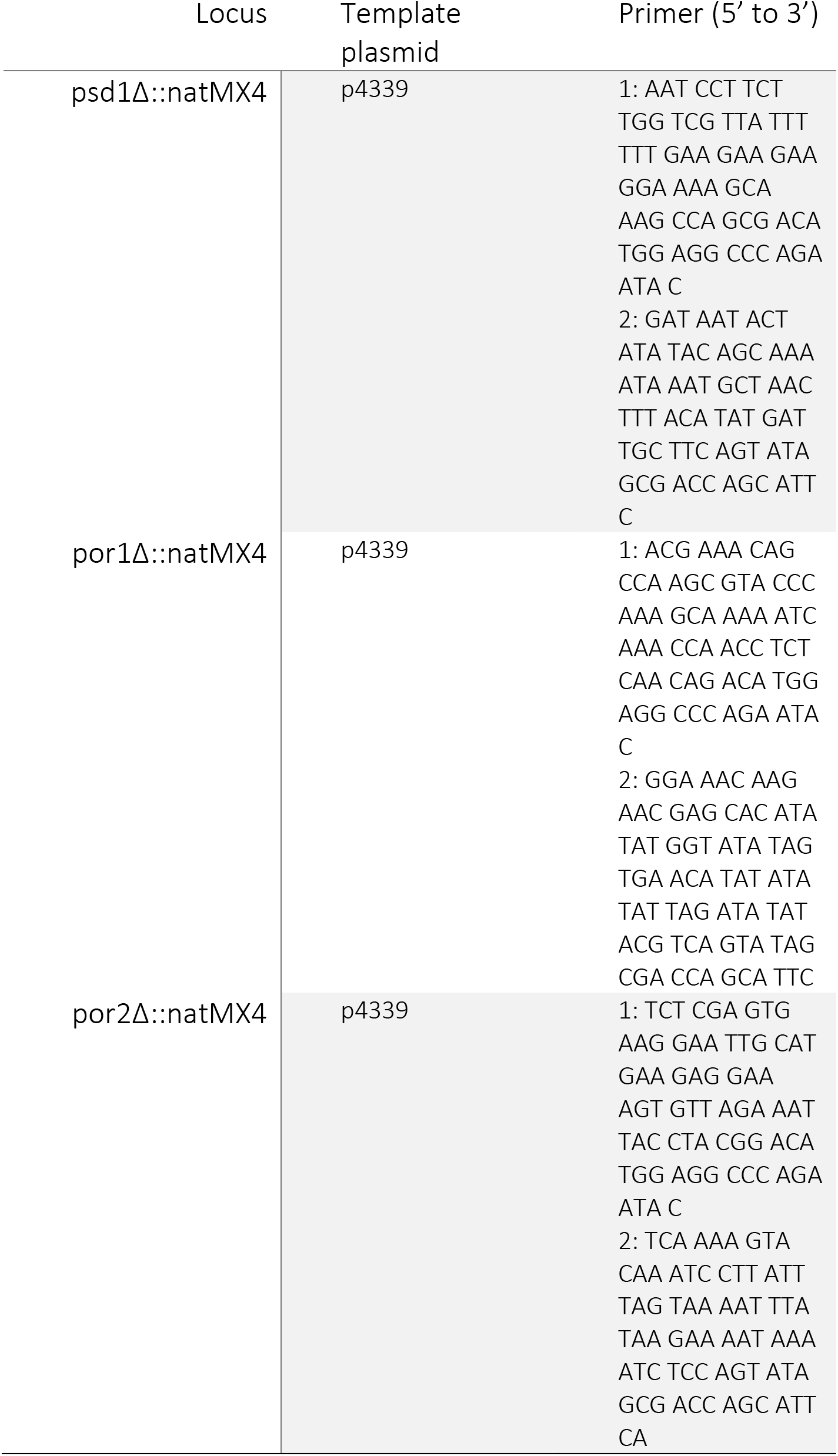
PCR templates and primers used in this study.

The initial structure for the VDAC1-5V mutant was generated by taking the structure of wild-type VDAC1 monomer and replacing residues T77, S43, T33, S35, Y247, and Q249 with valines using MODELLER version 9.11 (Sali & Blundell, 1993). The mutated monomeric structure was then minimized in vacuum, coarse-grained and a dimer-1 version was prepared in the same way as described above. Equilibration and production simulations and analysis were performed in the same way as for wild-type dimer-1 (described above).

## ACKNOWLEDGEMENTS

We thank Steve Claypool (Johns Hopkins University) for antibodies, Non Miyata (Kyushu University) for yeast strains, Matthew Johnson and Denisse Leyton (Australian National University) for the Pet464 expression construct, Kathryn Diederichs and Susan Buchanan (NIDDK) for the Por1 expression construct, and Toon de Kroon (Utrecht University) for the mitochondria isolation protocol. This work was supported by National Institutes of Health grants NS119779 (AKM) and NS116747 (JSD), Boehringer Ingelheim Fonds Ph.D. Fellowship (HJ), Deutsche Forschungsgemeinschaft grants SFB944-P14 and HO 3539/2-1 (JH), Czech Science Foundation grant GA20-20152S (RV), European Research Council (ERC) under the European Union’s Horizon 2020 research and innovation programme (grant agreement No 101001470) (RV), and National Institute of virology and bacteriology (Programme EXCELES, ID Project No. LX22NPO5103) - Funded by the European Union - Next Generation EU (RV). Computational resources were provided by the CESNET LM2015042 and the CERIT Scientific Cloud LM2015085 under the program Projects of Large Research, Development, and Innovations Infrastructures. Additional computational resources were obtained from IT4 Innovations National Supercomputing Center -- LM2015070 project supported by MEYS CR from the Large Infrastructures for Research, Experimental Development, and Innovations.

## AUTHOR CONTRIBUTIONS

Conceptualization: HJ, JH, JSD, RV, AKM

Methodology: HJ, LB, GID, JSD, RV, AKM

Investigation: HJ, LB, GID

Visualization: HJ, LB, RV, AKM

Funding acquisition: HJ, JSD, JH, RV, AKM

Project administration: RV, AKM

Supervision: RV, AKM

Writing – original draft: HJ, LB, AKM

Writing – review & editing: HJ, LB, GID, JSD, JH, RV, AKM

## DATA AND MATERIALS AVAILABILITY

All data are available in the main text or the supplementary materials.

## SUPPLEMENTARY FIGURES S1-S12

**Figure S1.**
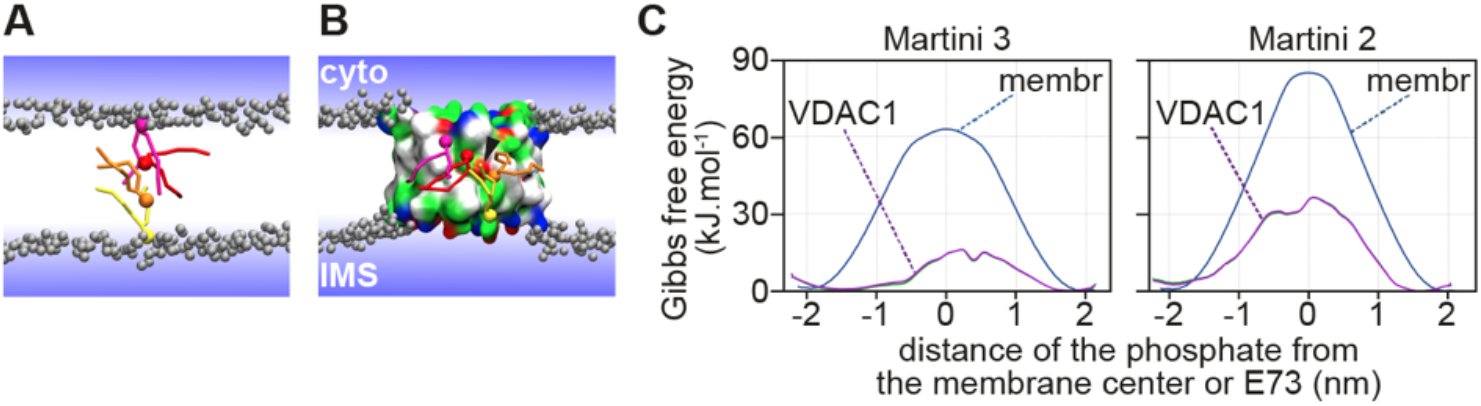
MD simulations to assess the energetics of phospholipid translocation in the presence of human VDAC1. **A, B.** Transbilayer phospholipid translocation was analyzed by coarse-grained molecular dynamics simulations of a pure POPC bilayer and a POPC bilayer containing human VDAC1 (PDB ID 6G6U). A phospholipid was pulled from one leaflet of the bilayer to the other in the z-direction. In the case of VDAC1, the lipid phosphate was also restrained to be within a cylinder, centered on E73, with a radius of 1.5 nm. Five selected positions along the translocation pathway from the upper (IMS) to lower (cytoplasmic) side are shown colored from yellow to red (the lipid headgroup is represented by single bead, and acyl chains are represented as thin tubes). Membrane phospholipids are not shown explicitly - their phosphate groups are shown by grey beads, while the lipid tails are not displayed for clarity. VDAC1 is represented as a protein surface color-coded as follows: hydrophobic-white, polar-green, red-negatively charged, and blue-positively charged residues. **C.** Calculated free energy profiles of lipid translocation as a function of the distance between the transiting lipid phosphate and the membrane center or E73 are shown. The presence of VDAC1 leads to a significant decrease in the energy barrier for phospholipid translocation around E73 using coarse-grained Martini 3 (left) or Martini 2 (right) force-fields. The error is estimated < 2 kJ.mol^-1^ for the pure POPC bilayer and < 5 kJ.mol^-1^ for the system with VDAC based on the profile asymmetry.

**Figure S2.**
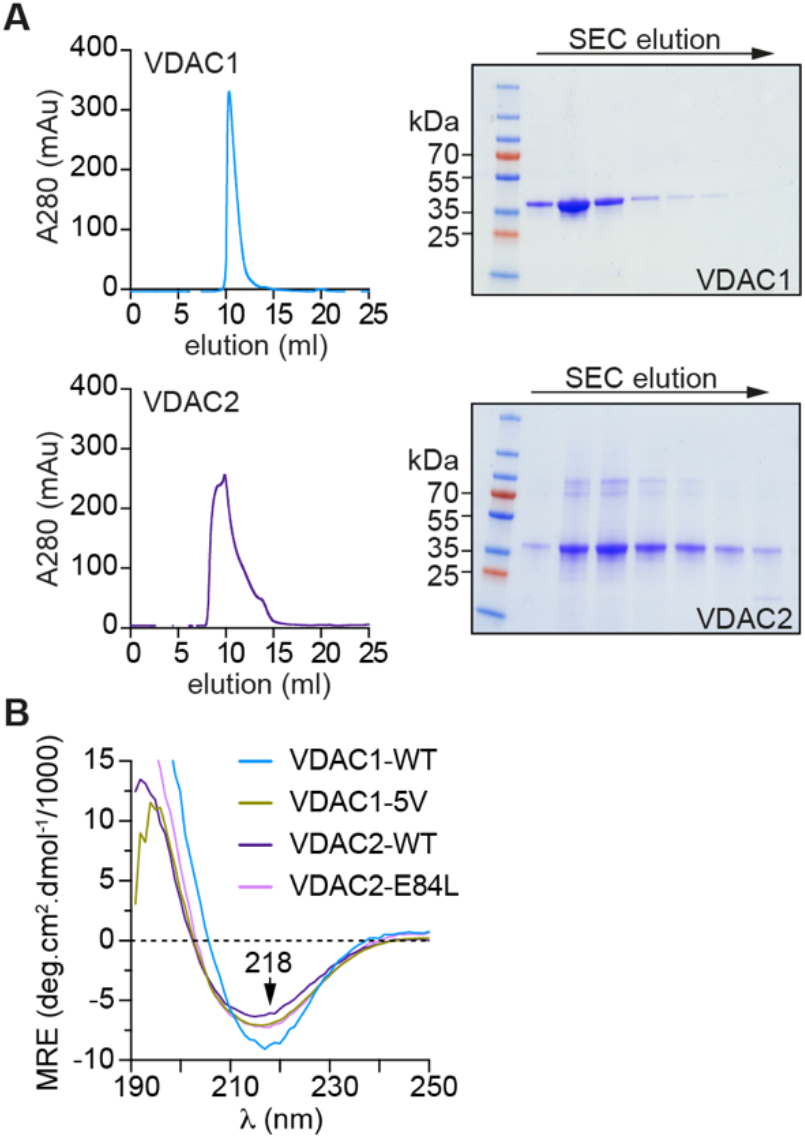
Purification of human VDAC1 and VDAC2. **A.** Human VDAC proteins were produced heterologously in *E. coli*, purified as inclusion bodies, denatured with guanidine hydrochloride, solubilized using the detergent *N*,*N*-Dimethyldodecylamine *N*-oxide (LDAO) and purified by cation-exchange and size-exclusion chromatography (SEC). SEC profiles and associated Coomassie-stained SDS-PAGE analyses are shown. **B.** Circular Dichroism spectroscopy of human VDAC proteins. Spectra were obtained with 66 µM VDAC samples (wild-type (WT) VDAC1 and VDAC2, VDAC1-5V mutant, and VDAC2-E84L mutant) in a 0.2 mm sandwich cell. All proteins showed high beta-sheet content.

**Figure S3.**
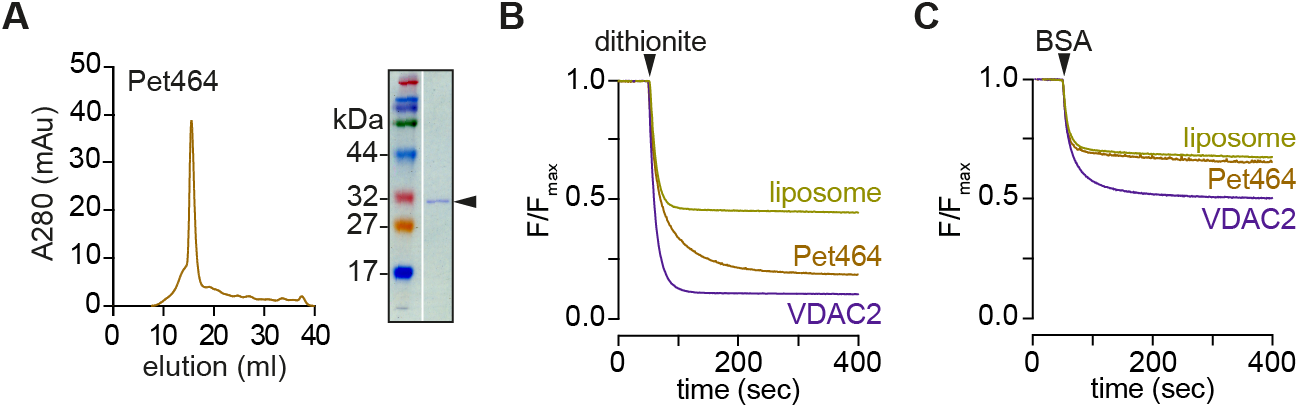
The Pet464 β-barrel protein does not scramble phospholipids. **A.** Size-exclusion chromatography (SEC) profile of Pet464, the β-barrel portion of the Pet autotransporter. The β-barrel was produced in *E. coli*, recovered as inclusion bodies, denatured with urea and re-folded in LDAO detergent. The refolded protein was purified using Ni-NTA affinity chromatography and SEC. The SEC profile and Coomassie-stained SDS-PAGE analysis of the peak fraction are shown; the arrowhead indicates the ∼30 kDa purified protein. **B, C.** Dithionite (B) and BSA-back extraction (C) assay traces comparing Pet464 with VDAC2 in vesicles containing NBD-PC as the reporter lipid. Both proteins were reconstituted at a predicted density of 60 copies per vesicle. Normalized fluorescence traces are shown.

**Figure S4.**
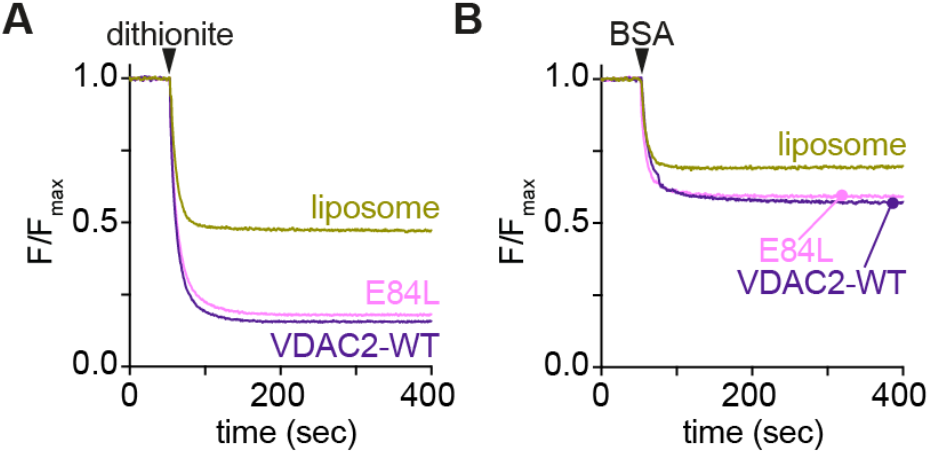
VDAC2 E84L is a scramblase. VDAC2-E84L was purified as in Figure S2 (CD spectrum shown in Fig. S2B), reconstituted into unilamellar vesicles containing NBD-PC at a theoretical protein/vesicle copy number of 10, and assayed for channel and scramblase activity. Wild-type VDAC2 was analyzed in parallel. **A.** Channel assay, as described in Fig. 1D. **B.** Scramblase assay, as described in Fig. 1F.

**Figure S5.**
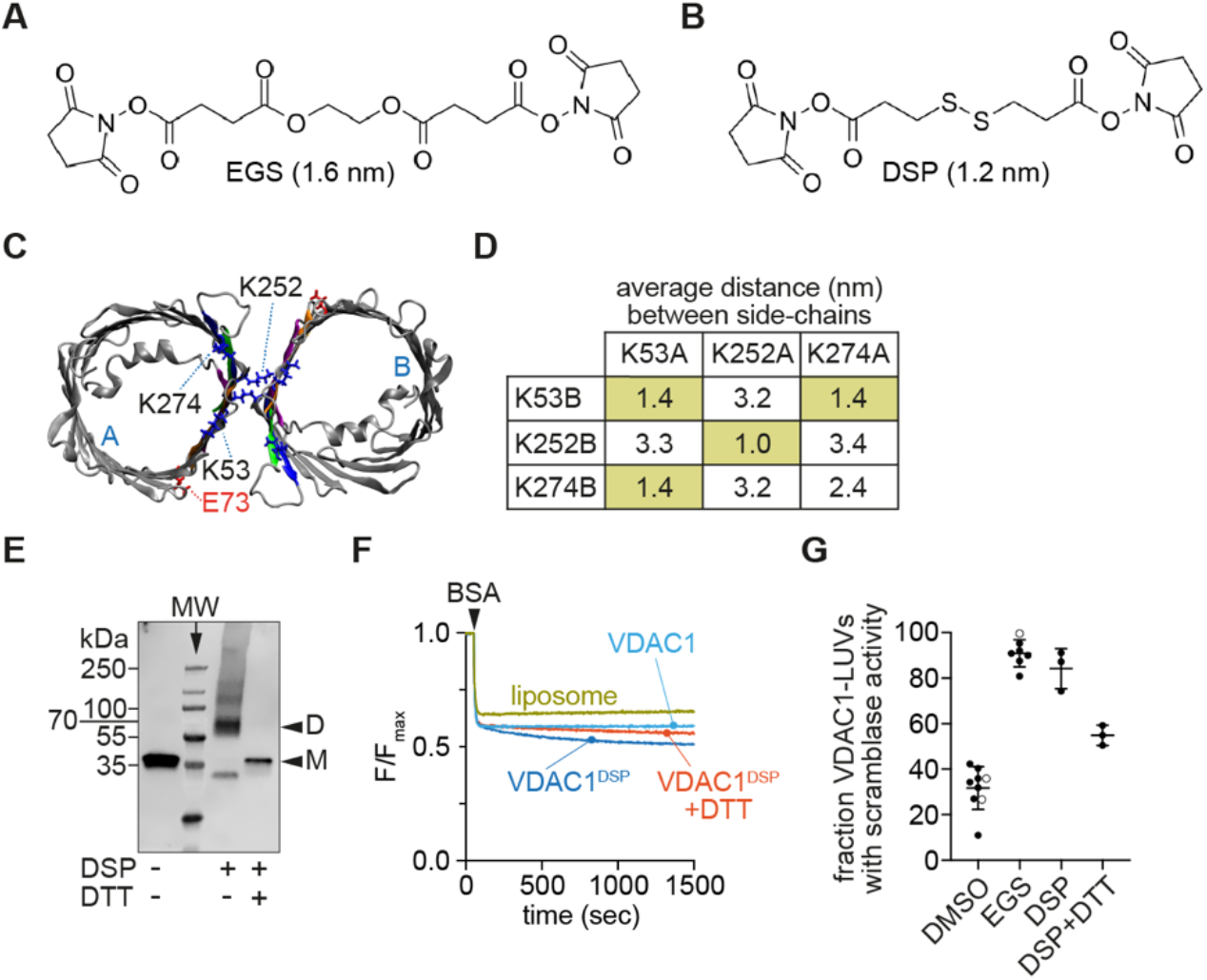
Scramblase activity of VDAC1 dimers prepared with a cleavable crosslinker. **A.** Chemical structure of EGS. **B.** Chemical structure of DSP. **C.** Top view of dimer-1, indicating outward-facing lysine residues in protomers designated A and B. **D.** Distances between lysine pairs on VDAC1 protomers in dimer-1 calculated from a coarse-grained MD simulation that accounts for reorientation of the sidechains. Each of the indicated distances is the average xyz distance between the sidechain bead of lysines on protomers A and B. The shaded boxes indicate lysine pairs that are <1.4 nm apart and therefore likely to be crosslinked by EGS and DSP. **E.** VDAC1 in LDAO was cross-linked using DSP or mocked treated with DMSO (indicated (-)) and visualized by SDS-PAGE immunoblotting using antibodies against the N-terminal His tag. An aliquot of the DSP-cross-linked sample was treated with dithiothreitol (DTT) after reconstitution into liposomes and analyzed alongside. **F.** Cross-linked (VDAC^DSP^) or mock-treated (VDAC1) proteins were reconstituted into LUVs at a copy number of 30 proteins/vesicle and assayed for scramblase activity using BSA back extraction as in Fig. 1F. Normalized fluorescence traces are shown. Also shown is an assay in which a VDAC^DSP^-reconstituted sample was treated with DTT to reverse the crosslink *in situ*. **G.** Fraction of VDAC1-vesicles with scramblase activity; data shown correspond to reconstituted DMSO-treated VDAC1, reconstituted EGS-or DSP-crosslinked VDAC1, and VDAC1^DSP^ treated with DTT after reconstitution at a theoretical protein/vesicle copy number of 30. Open circles in the DMSO and EGS data represent samples that were treated with DTT as a control.

**Figure S6.**
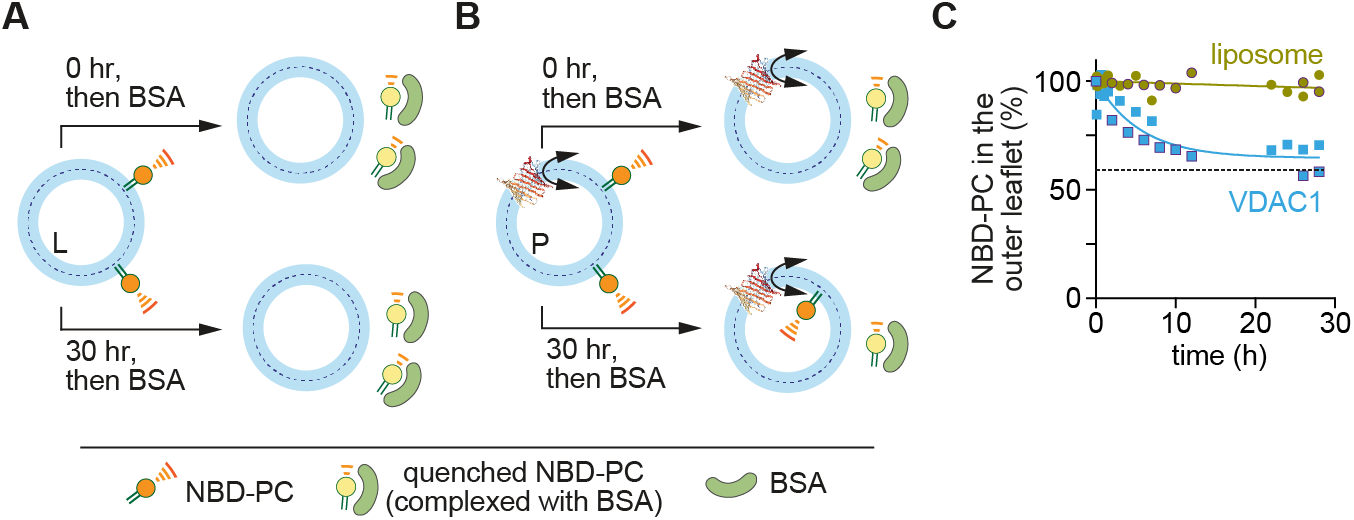
VDAC1 monomers scramble phospholipids slowly. **A, B.** Schematic of assay to detect slow scrambling. Asymmetric liposomes were prepared by adding 0.25 mol% NBD-PC lipids in EtOH (final EtOH concentration 0.4%) to protein-free liposomes (L) (panel A) or VDAC1-containing proteoliposomes (P) reconstituted at a theoretical protein/vesicle copy number of 10 (panel B). Samples were incubated for 30 min on ice after which they were incubated for approximately 30 h at 20°C. Aliquots were taken at different time points to perform a BSA-back extraction assay and determine the fraction of NBD-lipids present in the outer leaflet. If a slow scramblase is present, over the time course of 30 h, approximately 50% of NBD lipids should be shielded from BSA extraction at the inner leaflet. **C.** Slow scrambling by VDAC1. Two independent experiments with VDAC1-proteoliposomes are shown, fitted to a single exponential decay function. Parallel experiments with protein-free liposomes are also shown. The black dotted line indicates the expected fluorescence signal for the VDAC1-proteoliposomes, experimentally determined from protein reconstitutions with same conditions. The entire complement of NBD-PC molecules in the protein-free vesicles was accessible for BSA-extraction over the time-course of the experiment, indicating little or no scrambling. In contrast, for vesicles reconstituted with hVDAC1, nearly 50% of the NBD-lipids were shielded from BSA-extraction after 20 h, indicating a scrambling half-life ∼4 h.

**Figure S7.**
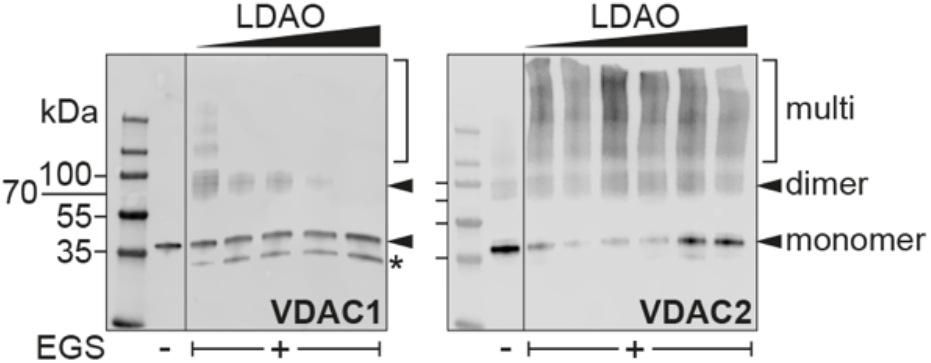
Crosslinking of LDAO-solubilized VDAC proteins. The oligomeric state of VDAC1 (left panel) and VDAC2 (right panel) in different concentrations of LDAO (0.05% - 1.05% w/v) was determined by cross-linking using the homobifunctional, amine-reactive crosslinker EGS at a 16-fold molar excess over lysine residues in the protein. EGS was delivered from a stock solution in DMSO (final DMSO concentration = 0.5%) and the reaction was carried out for 40 min at 20°C. Control samples (indicated (-)) were mock-treated with DMSO. The crosslinking reactions were analyzed by SDS-PAGE immunoblotting with anti-His tag antibodies. Oligomerization is highly dependent on the detergent concentration with mainly (VDAC1) or a greater proportion (VDAC2) of monomers present at high detergent concentrations.

**Figure S8.**
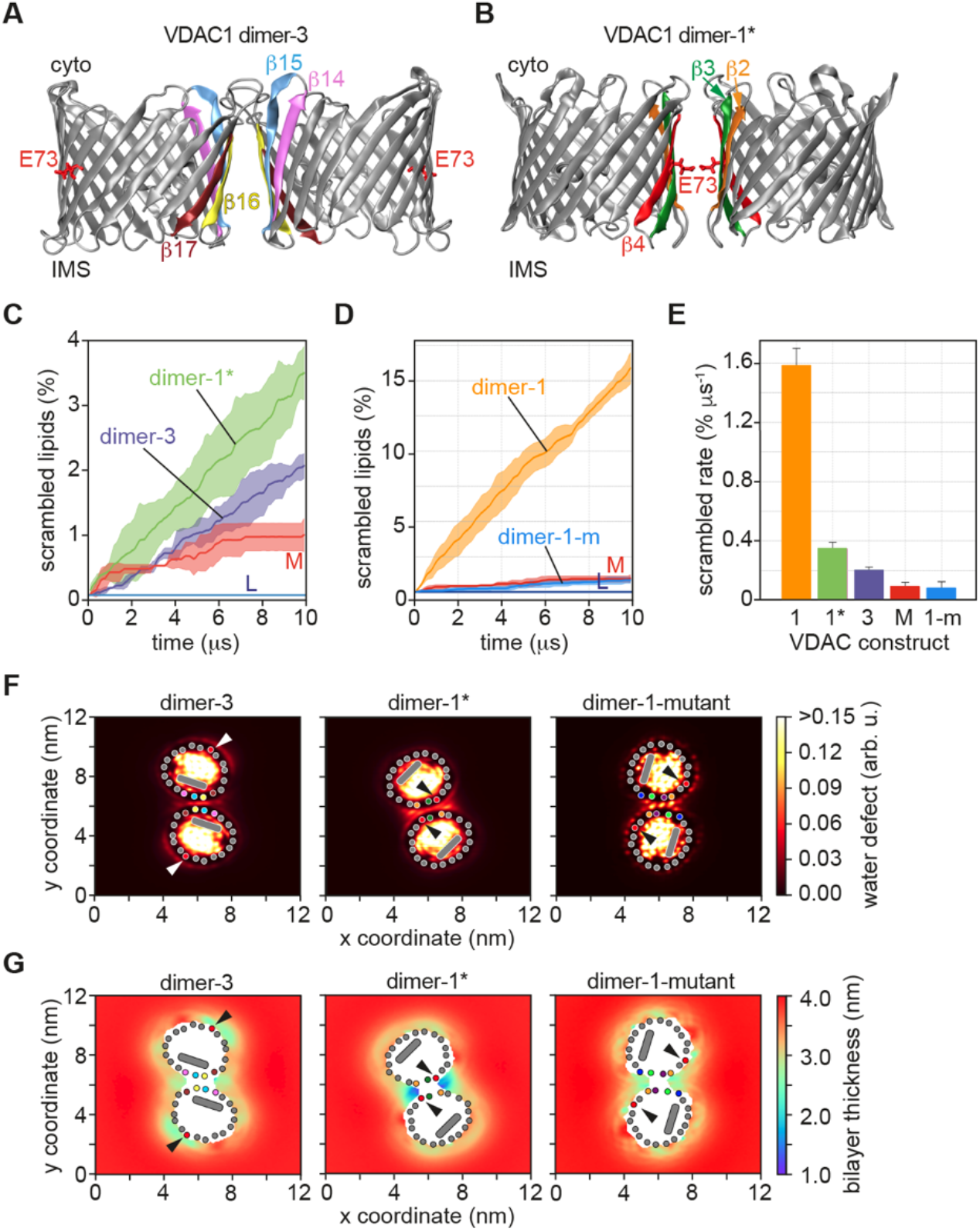
Structures of various VDAC1 dimers and their effects on the membrane. **A.** Structure of VDAC1 dimer-3. The β-strands at the interface are colored as indicated, and the E73 residue on the β4-strand is marked. **B.** Structure of VDAC1 dimer-1*; details are as in panel A. **C.** Percentage of lipids scrambled as a function of simulation time. The graphs show the scrambling activity of dimer-1*, dimer-3, monomer (M), and protein-free membrane (L) (running average over 200 ns time intervals, shading=68% confidence interval, n=3). **D.** As in panel C for constrained dimer-1 (taken from Fig. 4C) and constrained dimer-1-mutant (dimer-1-m). The mutations in dimer-1-m are T77V, S43V, T33V, S35V, Y247V, Q249V. **E.** Bar chart showing the scrambling rate for dimer-1, -1*, -3, monomer (M) (taken from Fig. 4D), and dimer-1-m (mean ± SD, n=3). **F.** Top views of water penetration into the membrane (water defect, color LUT shown at right) in the vicinity of the dimer interfaces. The β-strands at the interface are indicated as colored dots (same color scheme as in panels A, B); the E73 residue is shown as a red dot and marked by arrowheads; the N-terminal helix is shown as a grey oblong within the VDAC1 pore. **G.** Bilayer thinning at the dimer interfaces (thickness scale, color LUT shown at right, E73 indicated by arrowhead). Other details are as in panel F.

**Figure S9.**
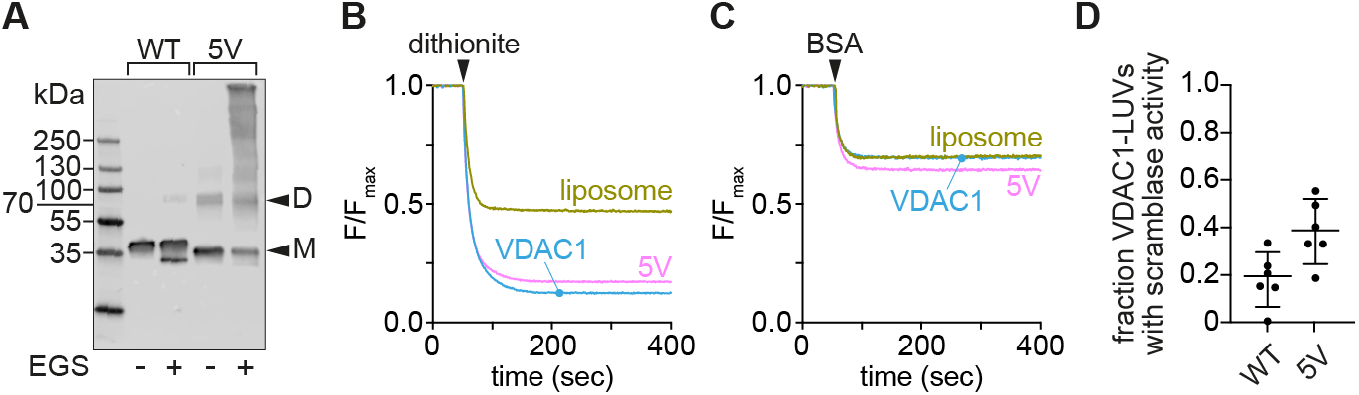
Analysis of the VDAC1-5V mutant. **A.** VDAC1-5V is a dimer/oligomer in liposomes. VDAC1 WT and the 5V mutant were reconstituted into unilamellar vesicles at a theoretical protein/vesicle copy number of 30 and subjected to EGS crosslinking to determine their oligomeric state. Control samples were mocked treated with DMSO. The samples were analyzed by SDS-PAGE immunoblotting using antibodies against the N-terminal His tag, revealing the 5V mutant to be highly dimeric/oligomeric. **B.** Representative channel assay as described in Fig. 1D of reconstituted VDAC1-WT and VDAC1-5V at a theoretical protein/vesicle copy number of 10. **C.** Representative scramblase assay as described in Fig. 1F of reconstituted VDAC1 WT and VDAC1-5V at a theoretical protein/vesicle copy number of 10. **D.** Fraction of VDAC1-vesicles with scramblase activity reconstituted at a theoretical protein/vesicle copy number of 10. Despite its dimeric/multimeric nature after reconstitution as evidenced by EGS crosslinking, the 5V mutant is not an effective scramblase.

**Figure S10.**
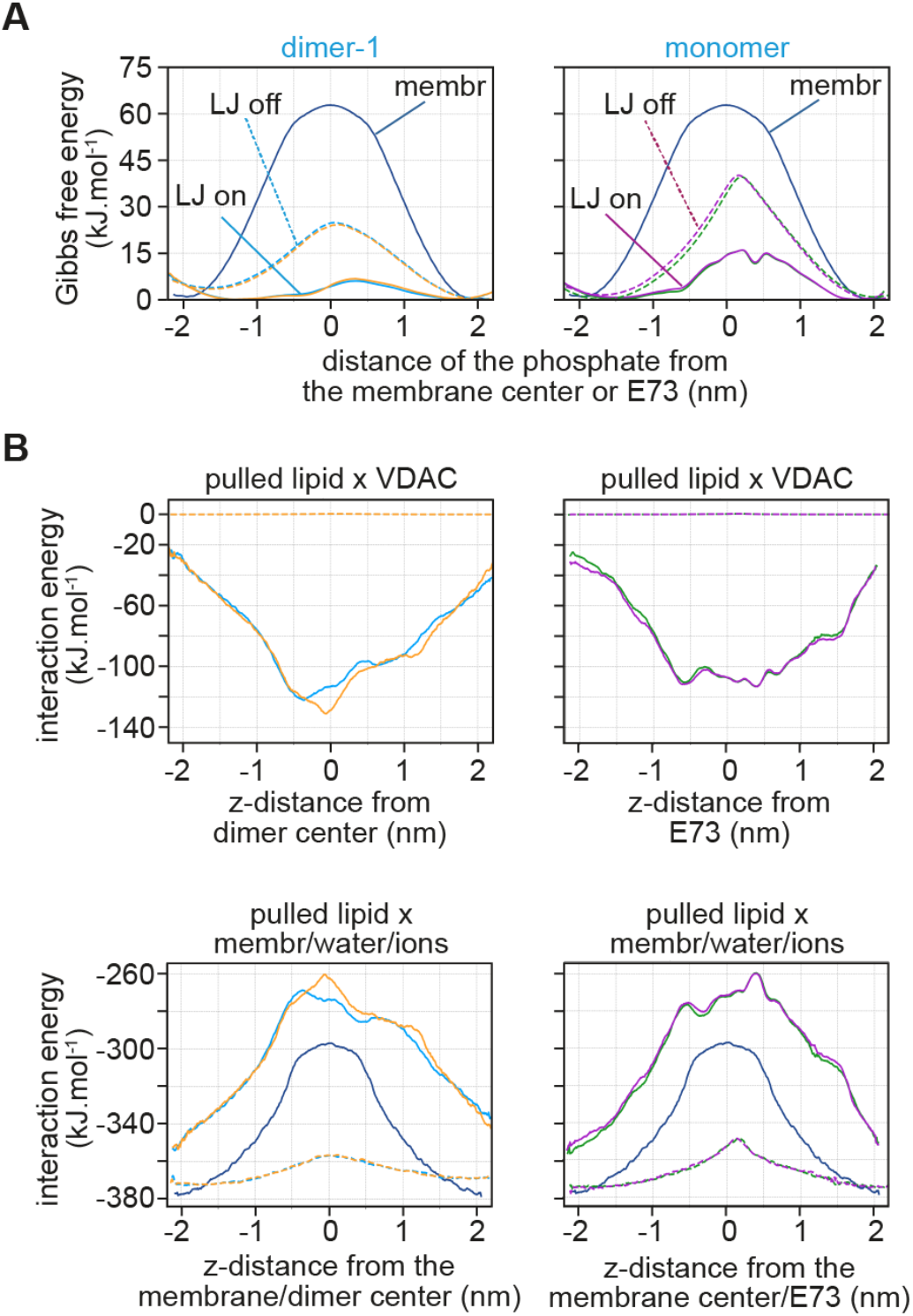
The effect of VDAC1 and membrane disruption on the energetics of lipid scrambling. **A.** Free energy profiles of phospholipid translocation as a distance between the lipid phosphate and the membrane center in pure POPC membrane (blue line), in the presence of VDAC1 monomer (green and magenta lines in the right panel), and in the presence of dimer-1 (azure and orange lines in the left panel). Dashed lines correspond to simulations in which the attractive dispersion interactions (Lennard-Jones (LJ) interactions) between the lipid and VDAC1 were turned off, effectively removing the direct contribution of VDAC1 to the decrease in the flip-flop barrier. **B,C.** Average enthalpic interactions (Lennard-Jones + electrostatic) between the scrambling lipid and VDAC (B) or between the scrambling lipid and membrane + water + ions (C) as a distance between the scrambling lipid phosphate and the membrane center. The enthalpic interactions were calculated from the simulated umbrella sampling windows. The color scheme is the same as in A.

**Figure S11.**
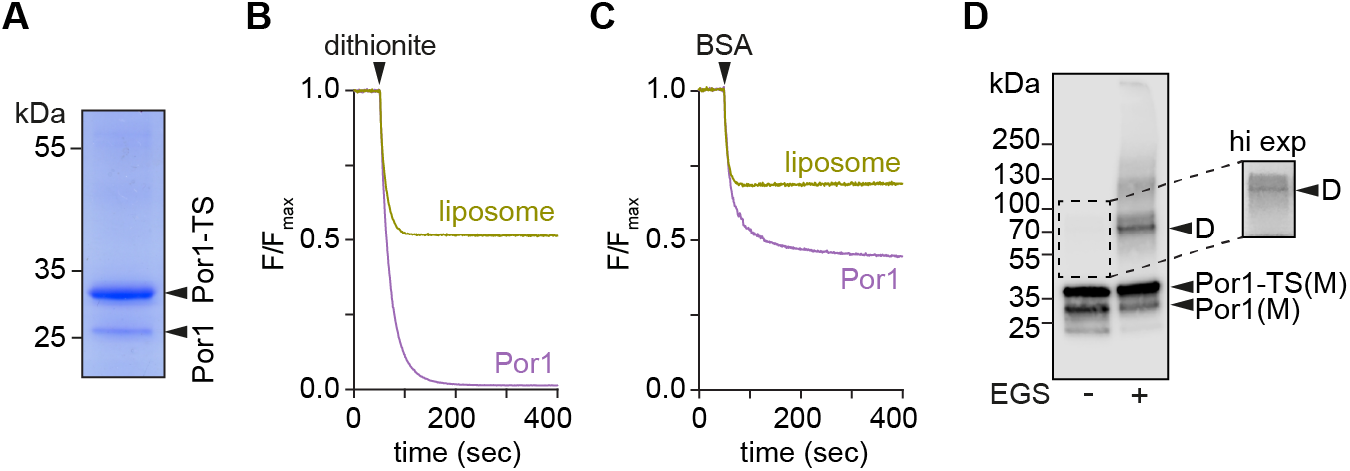
Native yeast VDAC (Por1) scrambles phospholipids. **A.** Coomassie-stained SDS-PAGE of Twin-Strep-tagged yeast VDAC (Por1-TS). The construct was expressed in W303 yeast cells, extracted from cell membranes in LDAO detergent, and purified on Strep-Tactin XT resin, followed by size-exclusion chromatography. Mass spectrometry confirmed that both bands on the gel correspond to Por1: the strongly stained upper band is Por1-TS, and the lower band is endogenous Por1, which copurifies with Por1-TS. The term ’native’ signifies that this protein was extracted from cells in folded form, in contrast to the human VDAC1 and VDAC2 preps (Fig. S2) where protein was recovered in inclusion bodies, denatured and re-folded. **B, C.** Normalized fluorescence traces corresponding to dithionite (B) and BSA-back extraction (C) assays performed on mock-treated liposomes or vesicles reconstituted with 12 ug of Por1-TS. Data points are shown as the mean of technical replicates (n = 3). **D.** Crosslinking of Por1 after reconstitution into vesicles. Reconstituted LUVs were treated with EGS crosslinker. The samples were analyzed by SDS-PAGE immunoblotting using anti-Por1 antibodies. Reconstituted Por1 shows a significant population of dimers/multimers on crosslinking. In the absence of EGS, high-exposure (hi-exp) of the blot reveals the presence of SD-resistant Por1 dimers.

**Figure S12.**
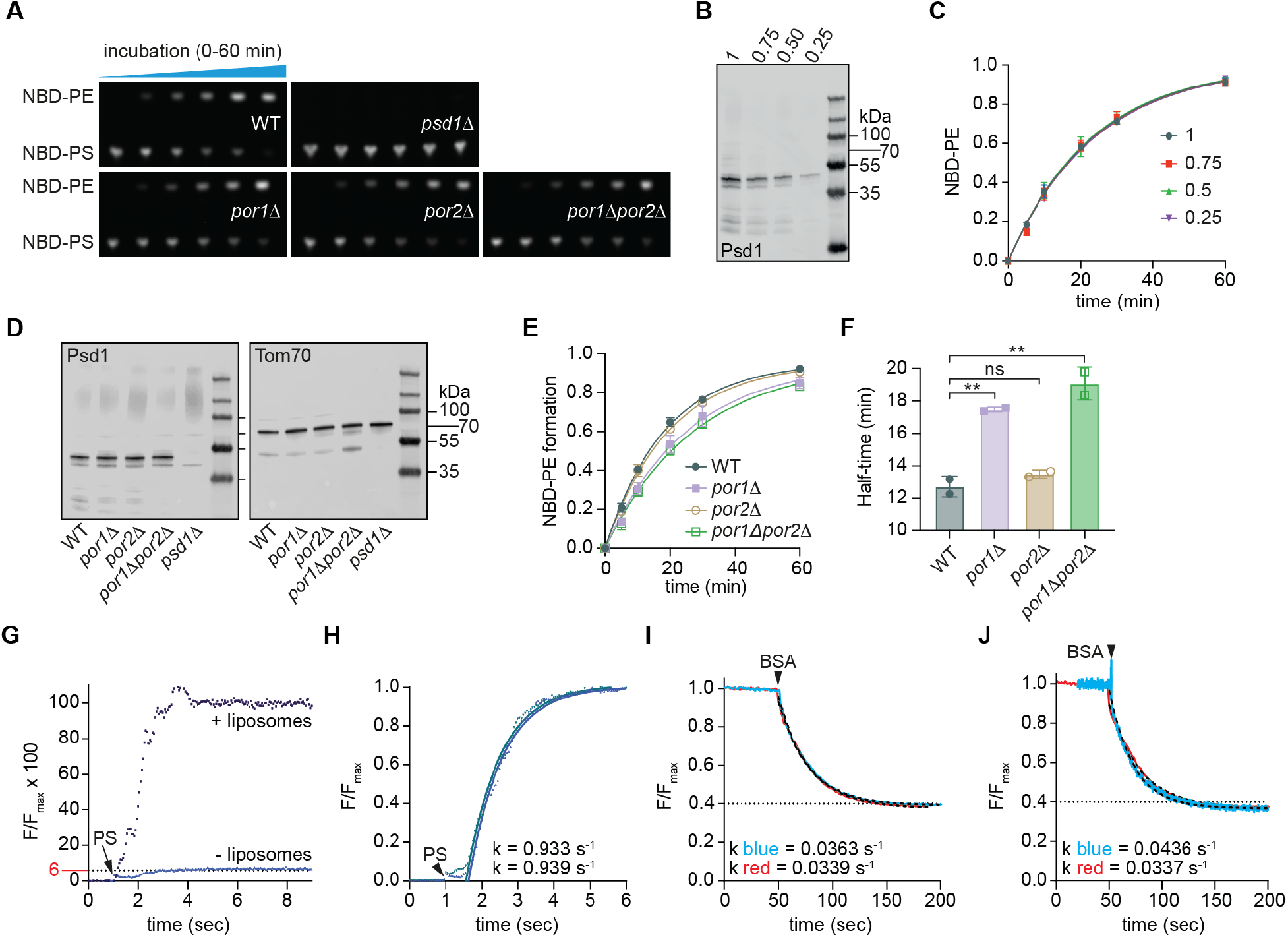
Supporting data for the PS-PE decarboxylation assay. **A.** Thin layer chromatograms, visualized with a ChemiDoc imager, of decarboxylation assay time courses using mitochondria from the indicated yeast strains. **B.** Different concentrations of mitochondria from wild-type cells (0.25-1 mg/ml, as indicated) were prepared for the decarboxylation assay. Equivalents were analyzed by immunoblotting using anti-Psd1 antibodies. **C.** Time-course of NBD-PS decarboxylation at 30°C using mitochondria at different concentrations as shown in panel C supplemented with 1 µM NBD-PS at time=0 min. The data points are shown as mean ± SD, fitted to a single exponential function. The rate of NBD-PE production was not affected by lowering the amount of mitochondrial protein 4-fold, confirming that mitochondria are not limiting in the assay. **D.** Mitochondria were isolated from the indicated yeast strains, and adjusted to have the same protein concentration, confirmed by immunoblotting using antibodies against the OMM protein Tom70 and the IMM protein Psd1. **E.** Time-course of NBD-PS decarboxylation at 30°C using mitochondria (1 mg/mL) from the indicated yeast strains, supplemented with 1 µM NBD-PS at time=0 min. The data points - obtained from chromatograms such as those shown in panel A - are given as mean ± SD and fitted to a single exponential function (2 biological replicates, with at least 2 technical replicates per assay). No NBD-PE was detected in *psd1Δ* cells (panel A). **F.** Half-time of NBD-PS to NBD-PE conversion obtained by mono-exponential fitting of the time traces shown in panel E (mean ± SD, p ≈ 0.0021 (One-Way ANOVA followed up with multiple comparison test, corrected using Bonferroni)). Fitting and analysis was carried out using GraphPad Prism. **G.** Fluorescence difference between NBD-PS monomer and NBD-PS in a POPC:POPG membrane, determined by adding 0.1 µM NBD-PS to liposomes (+liposomes) or buffer (-liposomes). After background subtraction, the traces were normalized to the maximum fluorescence (F_max_) of the +liposome sample. **H.** k_0_ was determined by adding 1 µM NBD-PS to POPC:POPG liposomes and fitting the fluorescence increase to a single exponential function. Two individual traces, and associated rate constants (k) are shown. **I.** BSA back extraction of asymmetrically labeled liposomes (1 µM NBD-PS total in sample) was performed to determine k_1_. Two traces are shown with different BSA concentrations (40 µl of 75 mg/ml BSA in red, or 40 µl of 37.5 mg/ml BSA in blue). The traces were analyzed using one-phase exponential decay; rate constants (k) are shown. **J.** BSA back extraction of asymmetrically labeled liposomes as in panel I, except that liposomes were prepared with either 1 µM NBD-PS (red trace) or 0.05 µM NBD-PS (blue trace), the latter to prevent micelle formation. Traces were analyzed as in panel I and the calculated rate constants are shown.

## MOVIE S1

**Movie S1. Phospholipid scrambling at the dimer-1 interface.**

Movie capturing spontaneous scrambling of a lipid along the interface of dimer-1 in a coarse-grained molecular dynamics simulation. The dimer is shown using molecular surface, the headgroups of most lipids are shown as gray beads and their tails as gray tubes, while a single lipid is highlighted in orange. Most lipids in front of the dimer are not depicted, except for several phosphate groups that are located close to the dimer. The movie captures roughly 140 ns of simulated time.

## SUPPLEMENTARY TABLES

